# Predator avoidance and foraging for food shape synchrony and response to perturbations in trophic metacommunities

**DOI:** 10.1101/2021.06.23.449565

**Authors:** Pierre Quévreux, Rémi Pigeault, Michel Loreau

## Abstract

The response of species to perturbations strongly depends on spatial aspects in populations connected by dispersal. Asynchronous fluctuations in biomass among populations lower the risk of simultaneous local extinctions and thus reduce the regional extinction risk. However, dispersal is often seen as passive diffusion that balances species abundance between distant patches, whereas ecological constraints, such as predator avoidance or foraging for food, trigger the movement of individuals. Here, we propose a model in which dispersal rates depend on the abundance of the species interacting with the dispersing species (*e.g.*, prey or predators) to determine how density-dependent dispersal shapes spatial synchrony in trophic metacommunities in response to stochastic perturbations. Thus, unlike those with passive dispersal, this model with density-dependent dispersal bypasses the classic vertical transmission of perturbations due to trophic interactions and deeply alters synchrony patterns. We show that the species with the highest coefficient of variation of biomass governs the dispersal rate of the dispersing species and determines the synchrony of its populations. In addition, we show that this mechanism can be modulated by the relative impact of each species on the growth rate of the dispersing species. Species affected by several constraints disperse to mitigate the strongest constraints (*e.g.*, predation), which does not necessarily experience the highest variations due to perturbations. Our approach can disentangle the joint effects of several factors implied in dispersal and provides a more accurate description of dispersal and its consequences on metacommunity dynamics.

## Introduction

Space is one of the major components of ecosystem functioning and must be taken into account to properly understand food web dynamics. The metacommunity concept, which considers a collection of patches that sustain communities of organisms able to disperse in other patches, provides a particularly useful framework for incorporating space in community and ecosystem ecology (Loreau et al., 2003; Leibold et al., 2004; Gross et al., 2020). This concept has led to a rich corpus of studies exploring the consequences of species dispersal on food web stability and the spatial synchrony of connected populations (Amarasekare, 2008; Leibold and Chase, 2017). However, these studies have not explored how density-dependent dispersal (*e.g.* dispersal depending on prey or predator abundance) shapes spatial synchrony in metacommunities affected by stochastic perturbations.

In the context of anthropogenic pressures, species synchrony is key to assessing community stability (Loreau and de Mazancourt, 2008), because species with synchronous populations have a higher biomass temporal variability at regional scale (Wang et al., 2015). In particular, synchronous populations can become extinct simultaneously, thus leading to regional extinction (Blasius et al., 1999). Although dispersal generally tends to synchronise populations by coupling their dynamics (Abbott, 2011; Fox et al., 2013), the dispersal of particular trophic levels can lead to asynchrony of populations of other trophic levels (Koelle and Vandermeer, 2005; Pedersen et al., 2016; Quévreux et al., 2021).

Most previous theoretical studies have considered dispersal as a passive diffusive process for the sake of simplicity. However, the density of other organisms should affect dispersal, thus altering the coupling between patches and patterns of synchrony. Organisms not only migrate from high density patches to low density patches but also respond to intra- and interspecific interactions (Fronhofer et al., 2015; Jacob et al., 2015). In particular, they move to forage for food and avoid predators (see the notion of landscape of fear developed by Laundré et al. (2001, 2010)), and their dispersal rates are functions of prey and predator densities. Density-dependent dispersal has been shown empirically across many taxa via experiments (Hauzy et al., 2007; Fronhofer et al., 2018; Harman et al., 2020), although its consequences on food web dynamics have only been explored by mathematical models.

Post et al. (2000) proposed a model in which a top predator feeds on two prey species belonging to two distinct food chains, and the predator’s for-aging effort depending on an *a priori* preference and the relative abundance of each prey. They found that asymmetric preference leads to dampened oscillations and thus to a stabilisation of the biomass dynamics. Following this avenue, McCann et al. (2005) and Rooney et al. (2006) showed that a wide predator spatial range and asymmetric productivity between the coupled food chains lead to a stabilisation of food webs (dampened oscillations or increased resilience). From prey’s perspective, Mchich et al. (2007) showed that the predator-dependent dispersal of prey stabilises the dynamics by promoting the existence of a stable equilibrium, while isolated predator-prey systems experience oscillations.

To our knowledge, Hauzy et al. (2010) are the only authors who extensively explored the effects of density-dependent dispersal on food web dynamics. They compared the effects of passive dispersal and three different types of density-dependent dispersal (Fig.1) in a predator-prey metacommunity with two patches, while other studies only focused on a single type of density-dependent dispersal. They found that density-dependent dispersal in both prey and predators tends to desynchronise the various populations and stabilise regional predator biomass dynamics while destabilising prey’s regional biomass dynamics because of the increased top-down control of prey by predators.

**Figure 1:**
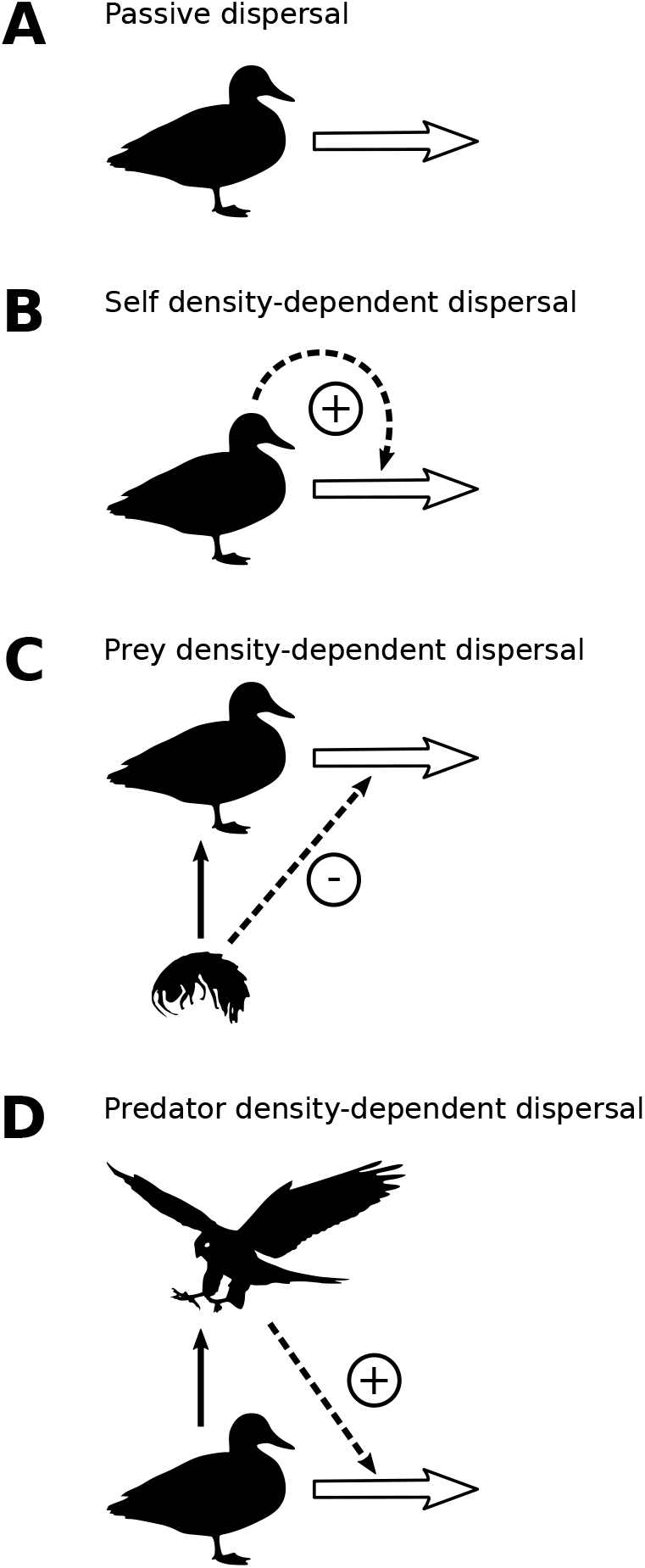
Different forms of dispersal. **A)** Passive dispersal, wherein dispersal rates are constant. Passive dispersal is similar to the passive diffusion of chemicals in the direction of a concentration gradient. **B)** Self density-dependent dispersal, wherein dispersal increases with the dispersing species biomass to avoid intraspecific negative interactions. **C)** Prey density-dependent dispersal, wherein dispersal decreases with prey biomass. **D)** Predator density-dependent dispersal, wherein dispersal increases with predator biomass.

Most theoretical studies on density-dependent dispersal define stability by the existence of equilibrium points or the amplitude of limit cycles (Li et al., 2005; Mchich et al., 2007; Hauzy et al., 2010; Liu et al., 2016). However, stability can also be defined by the response of the system to stochastic perturbations (Arnoldi et al., 2016, 2019), which is particularly relevant in the context of anthropogenic disturbances. Considering both limit cycles and stochastic perturbations is important be-cause different mechanisms affecting spatial synchrony are involved. Spatially correlated perturbations can synchronise distant populations (Moran effect Moran (1953)), and their transmission by particular trophic levels can lead to asynchrony due to trophic cascades (Quévreux et al., 2021). In the case of limit cycles, asynchrony is generally caused by phase locking, *i.e.* the coupling of the phases of the predator-prey oscillations in each patch (Jansen, 1999; Goldwyn and Hastings, 2008, 2009; Vasseur and Fox, 2009; Guichard et al., 2019).

A few studies considered the joint effects of stochastic perturbations and limit cycles. For instance, Vasseur and Fox (2007) showed that spatially correlated stochastic perturbations can dampen biomass oscillations but the effects of stochastic perturbations on metacommunities in the vicinity of equilibrium have only recently been studied (Wang et al., 2015; Jarillo et al., 2020; Quévreux et al., 2021). Our aim is to understand how density-dependent dispersal alters the synchrony and stability of populations in metacommunities in response to stochastic perturbations compared to previous studies that only considered passive dispersal.

### Box 1

#### Perturbation transmission in a metacommunity with passive dispersal

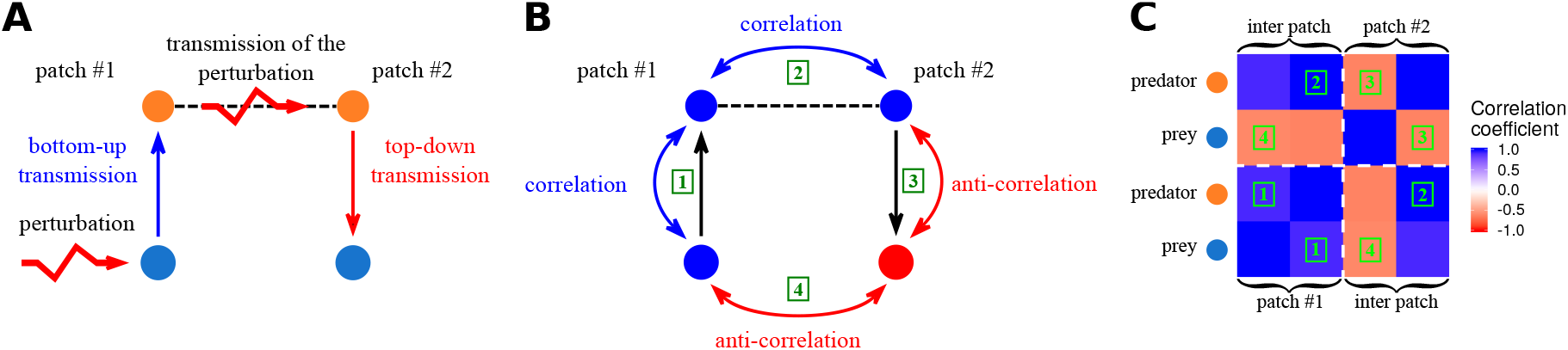

Here we summarise the main results from Quévreux et al. (2021), who considered a two patch predator-prey metacommunity with passive dispersal. In the setup presented in **A)**, prey are perturbed in patch #1 and only predators are able to disperse. Thus, perturbations have a bottom-up transmission in patch #1 (*i.e.* transmission from lower to upper trophic levels). This leads to the temporal correlation of the biomass dynamics of predators and prey in patch #1 showed in **B)** (1) because if a perturbation increases the biomass of prey, it also increases the biomass of predators due to the vertical transfer of biomass. The passive dispersal of predators transmits the perturbations and spatially correlate their populations as shown in **B)** (2). Then, perturbations have a top-down transmission in patch #2 (*i.e.* transmission from upper to lower trophic levels). This leads to the temporal anticorrelation (negative coefficient of correlation) of the biomass dynamics of predators and prey in patch #2 showed in **B)** (3) because if a perturbation increases the biomass of predators, it decreases the biomass of prey due to the negative effect of predators on prey. Eventually, prey populations are spatially anticorrelated, as shown in **B)** (4). Hence, by knowing which species is perturbed, which species disperses and how perturbations propagate within a food chain, Quévreux et al. (2021) were able to explain the spatial synchrony of the various populations of a metacommunity, summariesed by the correlation matrix in **C)**.

Here, we propose to extend the model developed in Quévreux et al. (2021), which describes a metacommunity with two patches sustaining Lotka-Volterra food chains, by integrating the density-dependent dispersal module proposed by Hauzy et al. (2010). Quévreux et al. (2021) were able to explain the spatial synchrony in multitrophic metacommunities affected by independent stochastic perturbations. They described how perturbations propagate vertically within patches and horizontally between patches, which leads an interpretation of spatial synchrony based on the trophic cascade framework used by Wollrab et al. (2012).

We expect density-dependent dispersal to modify these results. In fact, perturbations can be transmitted between patches without first being transmitted through trophic interactions because dispersal is directly affected by prey or predator density (Fig. 1), thus, the intrapatch vertical transmission of perturbations can be bypassed. While passive dispersal spatially correlates populations by balancing biomass distribution between patches (Fig. 1A), prey density-dependent dispersal (Fig.1C) should amplify the imbalance between patches by reducing the dispersal of consumers, whose biomass is growing due to prey abundance.

Then, we explore the joint effects of the various types of density-dependent dispersal described in Fig. 1. They are expected to have different relative importances since organisms have to balance foraging for food with predator avoidance to maximise their fitness (Gilliam and Fraser, 1987; Preisser et al., 2005; Preisser and Bolnick, 2008). For instance, predator density-dependent dispersal should contribute strongly to the overall dispersal rate of the prey if mortality due to predation is high. Considering realistic contributions of the various types of density-dependent dispersal should lead to a more accurate prediction of the responses of natural metacommunities to perturbations by accounting for the significant feedbacks of perturbation-driven changes in abundances on dispersal.

## Material and methods

### Metacommunity model

We use the model proposed by Quévreux et al. (2021) based on the food chain model developed by Barbier and Loreau (2019). The model consists of two patches that each sustain a food chain, and they are connected by species dispersal. We briefly present the food chain model in Box 2; however, a thorough description is provided in Barbier and Loreau (2019).

#### Box 2

##### Food chain model

The model has been originally developed by Barbier and Loreau (2019), who considered a food chain model with a simple metabolic parametrisation. Their model corresponds to the "intra-patch dynamics" part of equations (1a) and (1b) to which we graft a dispersal term to consider a metacommunity with two patches (see Box 1).

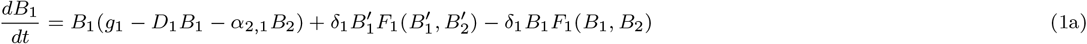

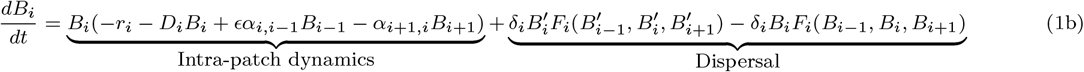

*B_i_* is the biomass of trophic level *i* in the patch of interest, 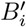 its biomass in the other patch, *ϵ* is the biomass conversion efficiency and *α_i,j_* is the interaction strength between consumer *i* and prey *j*. Species *i* disperses between the two patches at rate *δ_i_* and *F_i_*(*B*_*i*−1_, *B_i_*, *B_i+1_*) is the density-dependent dispersal function (see equation (5)). The density independent net growth rate of primary producers *g_i_* in equations (1a), the mortality rate of consumers *r_i_* in equations (1b) and the density dependent mortality rate *d_i_* scale with species metabolic rates *m*_*i*_ as biological rates are linked to energy expenditure.

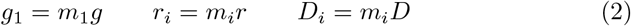

In order to get a broad range of possible responses, we assume the predator-prey metabolic rate ratio *m* and the interaction strength to self-regulation ratio *a* to be constant. These ratios capture the relations between parameters and trophic levels. This enables us to consider contrasting situations while keeping the model as simple as possible.

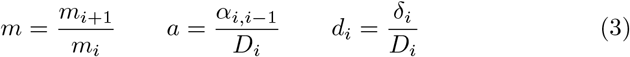

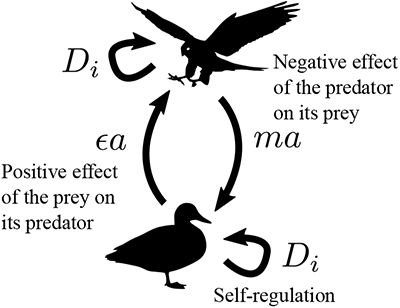

Varying *m* leads to food chains where predators have faster or slower biomass dynamics than their prey and varying *a* leads to food chains where interspecific interactions prevail or not compared with intraspecific interactions. As all biological rates are rescaled by *d_i_*, we also define *d_i_*, the dispersal rate relative to self-regulation (referred as scaled dispersal rate in the rest of the study), in order to keep the values of the dispersal rate relative to the other biological rates consistent across trophic levels. Finally, the time scale of the system is defined by setting the metabolic rate of the primary producer *m*_1_ to unity. Thus, we can transform equations (1a) and (1b) into:

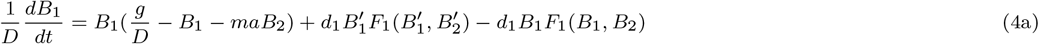

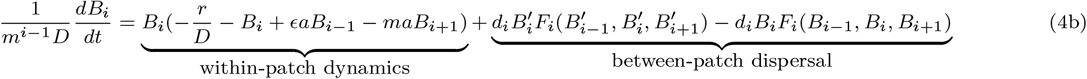

Thus, *ϵa* and *ma* defines the positive effect of the prey on its predator and the negative effect of the predator on its prey, respectively. These two synthetic parameters define the overall behaviour of the food chain and will be varied over the interval [0.1, 10] (see Table S1-1 in the supporting information) to consider a broad range of possible responses. Parameter values are summarised is Table 1.

**Table 1:** Table of parameters. Only combinations of *m* and *a* leading to the desired values of *ma* are retained.

| parameter | interpretation | value |
| --- | --- | --- |
| $\sigma_i$ | standard deviation of stochastic noise | $\{10^{-3}, 10^{-5}\}$ |
| $z$ | perturbation scaling exponent | $\{0.5, 1\}$ |
| $g$ | net growth rate of primary producers | 1 |
| $r$ | death rate of consumers | 0 |
| $D$ | self regulation | 1 |
| $\epsilon$ | conversion efficiency | 0.65 |
| $m$ | predator/prey metabolic rate ratio | $\{0.0065, 0.065, 0.65, 6.5, 65\}$ |
| $a$ | attack rate | $\{1/6.5, 1/0.65, 1/0.065\}$ |
| $\epsilon a$ | positive effect of prey on predators | $\{0.1, 1, 10\}$ |
| $ma$ | negative effect of predators on prey | $\{0.1, 1, 10\}$ |
| $d_i$ | scaled dispersal rate of species $i$ | $[10^{-5}, 10^5]$ |
| $C_{0,i}$ | passive dispersal component of species $i$ | $\{0, 1\}$ |
| $S_{0,i}$ | half saturation constant of density-dependent dispersal functions of species $i$ | $[10^{-4}, 10^4]$ |

Species dispersal depends on the abundance of the species living in the same patch. Following the empirical observations of Fronhofer et al. (2018), we consider that the density-dependent dispersal function *F_i_*(*B*_i-1_, *B*_*i*_, *B*_*i*+1_) (equation (5)), is the outcome of four additive components described in Fig. 1.

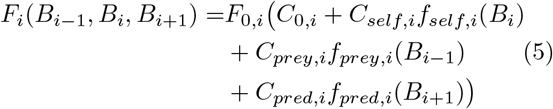

where *C*_0*,i*_ represents the classic passive dispersal if equal to 1; *f_self,i_*(*B_i_*) is the dependency of species *i* on its own density; *f_prey,i_*(*B*_*i*−1_) is the dependency on the density of prey; *f_pred,i_*(*B*_*i*−+1_) is the dependency on the density of predators; *C_self,i_*, *C_prey,i_* and *C_pred,i_* are the weights of each of these dependencies in the dispersal function; and *F*_0*,i*_ is a scaling constant set to obtain *F_i_*(*B*_*i*−1_, *B*_*i*_, *B*_*i*+1_) = 1 at equilibrium, as explained in the *Weighted dispersal components* subsection of the *Material and methods* section.

### Density-dependent dispersal

#### Dispersal component functions

Dispersal components *f_self,i_*(*B_i_*), *f_pred,i_*(*B*_*i*+1_) and *f_prey,i_*(*B*_*i*−1_) follow the functions defined by Hauzy et al. (2010) because of their interesting qualities. They are positive, which ensures dispersal flows are oriented in the right way, and symmetric, which facilitates the comparison between positive and negative density-dependent dispersal (Fig. 2). Components *f_self,i_*(*B_i_*) and *f_pred,i_*(*B*_*i*+1_) increase with *B_i_* and *B*_*i*+1_ (positive density-dependent dispersal) while *f_prey,i_*(*B*_*i*−1_) decreases with *B*_*i*−1_ (negative density-dependent dispersal).

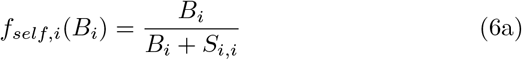

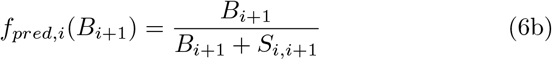

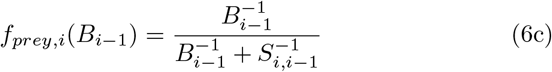

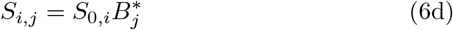

where *S_i,j_* is the sensitivity of the dispersal of species *i* to species *j* biomass density, with *S*_0*,i*_ a constant and 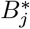 the biomass of species *j* at equilibrium. Such scaling ensures similar responses among species even if they have different biomasses.

**Figure 2:**
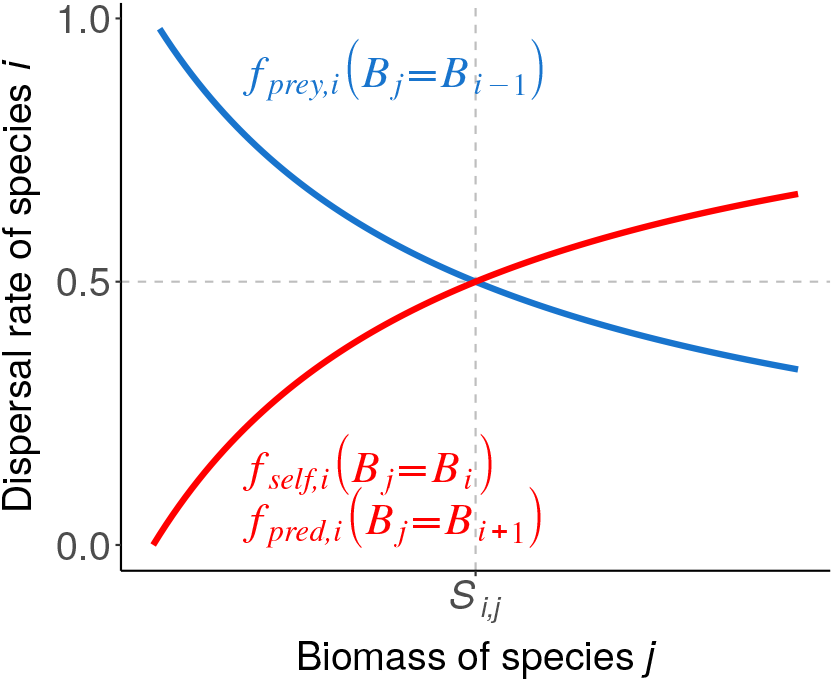
Functions describing the density-dependent dispersal rates of species *i* depending on species *j*. *B_j_/*(*B_j_* + *S_i,j_*) expresses self- and predator biomass dependency (red), and 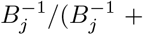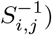 expresses prey biomass dependency (blue).

#### Weighted dispersal components

Here, the major challenge is to compare scenarios consisting of different combinations of the various types of density-dependent dispersal while keeping the overall dispersal rate constant. In other words, we aim to not change how many individuals disperse. Since we have no prior information on the values of dispersal rates among species, we rescale *F_i_*(*B_i-1_*, *B_i_*, *B*_*i*+1_) to keep its value equal to 1 by tuning the value of *F*_0*,i*_. Thus, only the scaled dispersal rate *d_i_* controls the intensity of dispersal, which makes correlation patterns comparable to the previous results of Quévreux et al. (2021).

Another major question is to determine the relative strength of the different components defined by *C_self,i_*, *C_prey,i_* and *C_pred,i_*. The null hypothesis is an equal strength, which is why term (1) in equation (7) rescales *C_self,i_*, *C_prey,i_* and *C_pred,i_* to 1. In nature, organisms tend to maximise their fitness by foraging for food and avoiding predators, which ultimately affect dispersal. For instance, predator density-dependent dispersal must contribute strongly to the overall dispersal rate if mortality due to predation is high. Hence, each dispersal component can be weighted by the associated within-patch process at equilibrium described in equations (4a) and (4b). *f_self,i_*(*B_i_*) can be weighted by the density-dependent mortality term 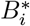, *f_prey,i_*(*B_i-1_*) by the prey consumption term 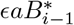 and *f_pred,i_*(*B*_*i*+1_) by the predation term 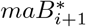 (term (2) in equation (7)).

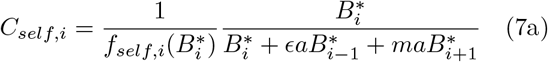

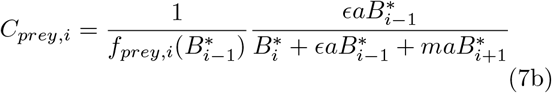

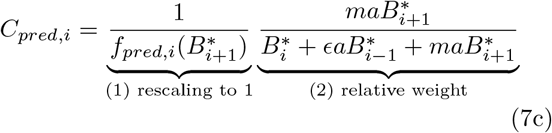

##### Box 3

###### Dispersal versus local demographic processes

Quévreux et al. (2021) demonstrated that the capacity of dispersal to couple populations and transmit perturbations highly depends on the importance of dispersal relative to local demographic processes (*e.g.* self-regulation and predation). They defined a metric *M*_1_ that describes this relative importance by taking the absolute values of the elements of equations (4a) and (4b) to assess the sheer intensity of local demographic processes and dispersal processes calculated with the equilibrium biomasses:

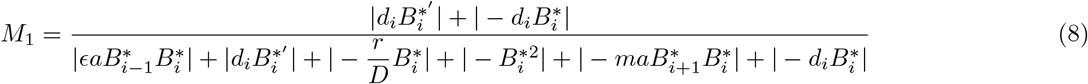

*M*_1_ is higher for less abundant species because dispersal scales linearly with biomass while intrapatch demography scales with squared biomass (self-regulation) or biomass products (predation) (see equations (4a) and (4b)).

As passive dispersal and self density-dependent dispersal lead to the same correlation patterns (see Fig. S2-1 and Fig. S2-2 in the supporting information), we consider *C*_0*,i*_ = 0 for the sake of simplicity.

### Stochastic perturbations

We use the same methods as Quévreux et al. (2021) to study the response of metacommunities to stochastic perturbations, and we provide only a brief description of the main concepts; thus, we invite readers to refer to their method section and supporting information S1 for a detailed description.

From equations (4a) and (4b), we obtain the following stochastic differential equation:

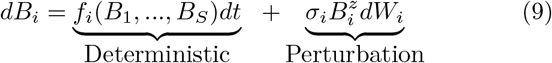

*f_i_*(*B*_1_, …, *B_S_*) represents the deterministic part of the dynamics of species *i* that depends on the biomass of the *S* species present in the metacommunity (as described by equations (4a) and (4b)). Stochastic perturbations are defined by their standard deviation *σ_i_* and *dW_i_*, a white noise term with a mean of 0 and variance of 1. In addition, perturbations scale with each species biomass with an exponent *z*. We consider two types of perturbations (Haegeman and Loreau, 2011; Arnoldi et al., 2019): demographic stochasticity (from birth-death processes) corresponds to *z* = 0.5 and environmental factors correspond to *z* = 1 (see demonstration in Lande et al. (2003) and in appendix S1-2 in the supporting information). We first consider demographic perturbations because Arnoldi et al. (2019) showed that they evenly affect each species regardless of abundance (*i.e.*, the ratio of species biomass variance to the perturbation variance is roughly independent of biomass), which enables us to describe the fundamental effects of density-dependent dispersal by perturbing specific species and avoiding any bias due to species abundance. Then, we consider environmental perturbations to study the response of our model in an ecologically relevant context.

### Response to perturbations

We aim to determine the synchrony between populations at equilibrium when they experience small stochastic perturbations. Synchrony can be evaluated from the covariances between the temporal variations of different species and patches, which are encoded in the variance-covariance matrix *C^∗^*. Therefore, we linearise the system in the vicinity of equilibrium to get equation (10) where 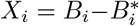 is the deviation from equilibrium (see section S1-3 and S2-19 in the supporting information).

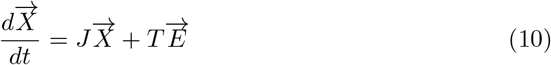

*J* is the Jacobian matrix (see section S1-4 in the supporting information) and *T* defines how the perturbations *E_i_* = *σ_i_dW_i_* apply to the system (scaling with species biomass).

We then obtain get the variance-covariance matrix *C^∗^* of species biomasses (variance-covariance matrix of 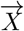) from the variance-covariance matrix of perturbations *V_E_* (variance-covariance matrix of 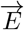) by solving the Lyapunov equation (11) (Arnold, 1974; Wang et al., 2015; Arnoldi et al., 2016; Shanafelt and Loreau, 2018).

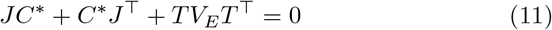

The expressions for *V_E_* and *T* and the method to solve the Lyapunov equation are detailed in section S2-19 in the supporting information. The variance-covariance matrix *C^∗^* can also be obtained through numerical simulations with the Euler-Maruyama method detailed in section S1-6 in the supporting information.

From the variance-covariance matrix *C^∗^* whose elements are *w_ij_* we can compute the correlation matrix *R^∗^* of the system whose elements *ρ_ij_* are defined as follows:

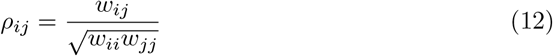

### Simulations design

In the following, we first consider food chains with two trophic levels, in which one trophic level is perturbed in patch #1 and only one trophic level is able to disperse, which enables us to explore the mechanisms of density-dependent dispersal. Then, we consider a setup with four tropic levels where all species in patch #1 are perturbed. We vary the scaled dispersal rate *d*_1_ and the sensitivity coefficient *S*_0*,i*_ to describe the response of spatial synchrony to the intensity of dispersal and the intensity of the density-dependency of dispersal.

## Results

### Density-dependent dispersal reverses spatial synchrony compared to passive dispersal

We start by describing the response of a simple metacommunity with two patches and a predatorprey system (Fig. 3). We consider two metacommunity setups to explore the effects of density-dependent dispersal: a setup where prey are perturbed in patch #1 and only predators are able to disperse (Fig. 3A and 3C) and a setup where predators are perturbed in patch #1 and only prey are able to disperse (Fig. 3B and 3D).

**Figure 3:**
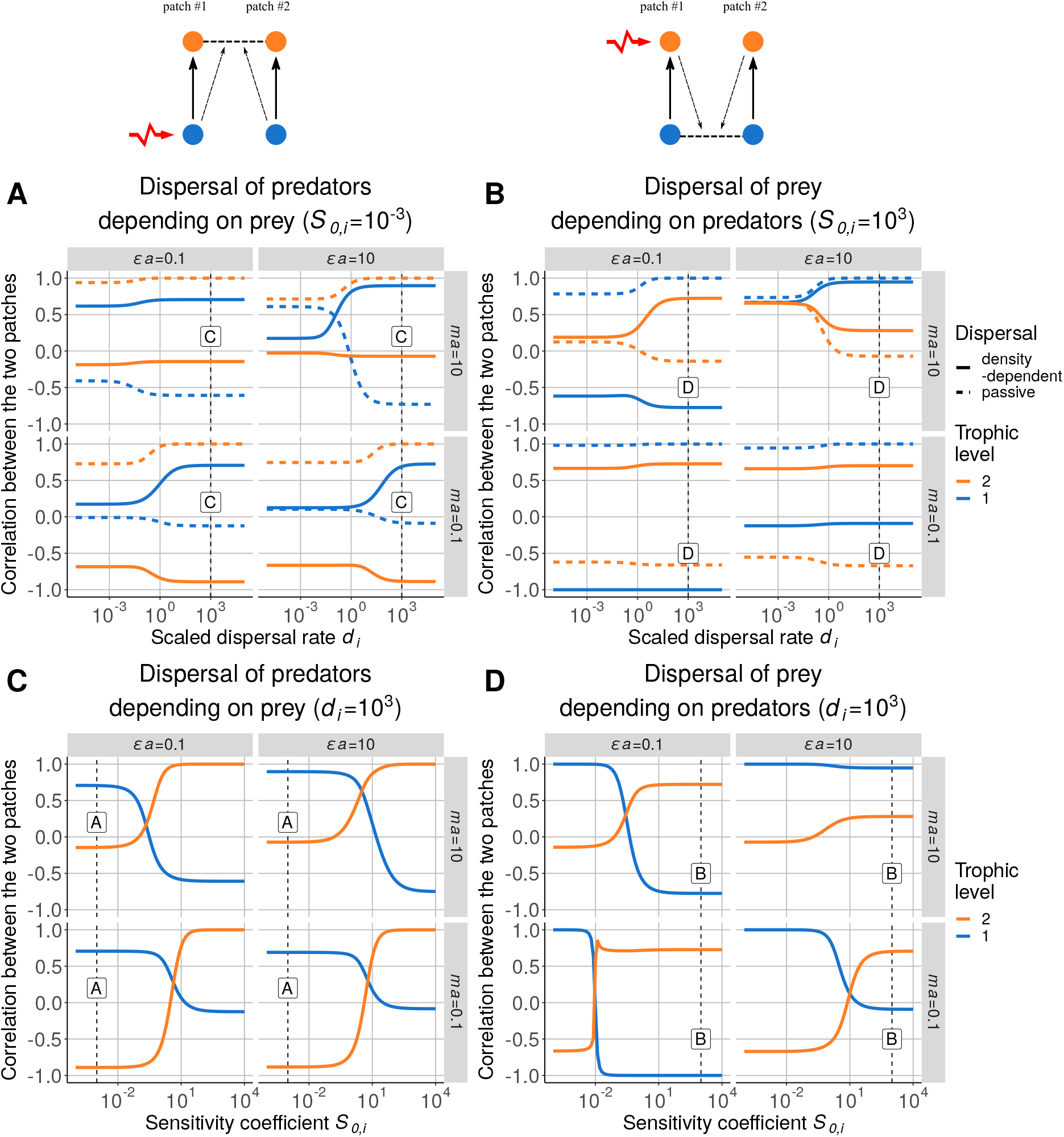
Correlation between the populations in each patch for increasing values of scaled dispersal rate *d_i_* when dispersal depends on **A)** the biomass of prey for dispersing predators (*S*_0*,i*_ = 10^−3^) and **B)** on the biomass of predators for dispersing prey (*S*_0*,i*_ = 10^3^). The dashed curves are the correlations for passive dispersal (reference from Quévreux et al. (2021)). Correlation for increasing values of sensitivity coefficient *S*_0*,i*_ (*d_i_* = 10^3^) when dispersal depends **C)** on the biomass of prey for dispersing predators and **D)** on the biomass of predators for dispersing prey. The dashed vertical lines represent the values of *d_i_* and *S*_0*,i*_ used in the other panels indicated by the labels overlapping the lines. *ϵa* is the positive effect of prey on predators and *ma* is the negative effect of predators on prey.

The results obtained with the model with passive dispersal are similar to that of Quévreux et al. (2021) and serve as a reference (dashed lines). In this case, for a high scaled dispersal rate *d_i_*, the two populations of the dispersing species are correlated while the populations of the nondispersing species tend to be anticorrelated (see Fig. 3A and 3B and Fig. S2-1 in the supporting information). In fact, perturbations have a bottom-up transmission in one patch, which correlates predators and prey dynamics, while they have a top-down transmission in the other patch, which anticorrelates the dynamics. Therefore, the two populations of the nondispersing species are anticorrelated. The mechanisms involved are thoroughly described in Quévreux et al. (2021) and are briefly described in Box 1.

Now, we consider that predator dispersal also depends on prey density in the setup where prey are perturbed in patch #1 and only predators are able to disperse (schema on top of Fig. 3A). In this case, the spatial synchrony is inverted compared to the case with passive dispersal for a sensitivity coefficient *S*_0*,i*_ = 10^−3^ (Fig. 3A), while it is similar for a sensitivity coefficient *S*_0*,i*_ = 10^3^ (see Fig. S2-1A and Fig. S2-3A in the supporting information). This inversion is visible on Fig. 3C where the value of *S*_0*,i*_ used in Fig. 3A is emphasised by the vertical dashed line and the letter A. Thus, for a sensitivity coefficient *S*_0*,i*_ = 10^−3^, if the density of prey increases in patch #1, then the density of predators in patch #1 increases because of the positive bottom-up effect and the decrease in the emigration rate. In patch #2, the density of predators decreases because emigration towards patch #1 is stronger than emigration towards patch #2 due to the lower abundance of prey in patch #2 (see the time series in Fig. S2-5 in the supporting information). Therefore, predator populations become anticorrelated.

Then, we consider that prey dispersal also depends on predator density in the setup where predators are perturbed in patch #1 and only prey are able to disperse (schema on top of Fig. 3B). In the same way, the spatial synchrony is inverted compared to the case with passive dispersal for a sensitivity coefficient *S*_0*,i*_ = 10^3^ (Fig. 3B) while it is similar for a sensitivity coefficient *S*_0*,i*_ = 10^−3^. Again, the inversion is clearly visible on Fig. 3D where the value of sensitivity coefficient *S*_0*,i*_ used in Fig. 3B is emphasised by the vertical dashed line and the letter B.

### Dispersal is driven by the species with highest biomass CV

However, we do not observe this inversion for negative effect of predators on prey *ma* = 10 and a positive effect of prey on predators *ϵa* = 10. For this set of parameters, the vertical biomass transfer is efficient and predators strongly deplete their prey (see Fig. S2-13A in the supporting information), which leads to a trophic cascade similar to those observed in pelagic food webs (Carpenter et al., 1985; Hulot et al., 2014). This absence of inversion is clearly visible on Fig. 3D (top-right panel). Depending on food chain length, species 1 (prey) and 2 (herbivore) are controlled or not by predation (Fig. 4A). Then, if species 1 is controlled (food chain length equal to 2 or 4), the two populations of species 1 remain correlated with increasing sensitivity coefficient *S*_0*,i*_ (Fig. 4B), while if species 1 is not controlled (food chain length equal to 3), they become anticorrelated. This switch of correlation is explained by the relative value of biomass CV between predators and prey. If the top-down control is weak (positive effect of prey on predators *ϵa* 1 and negative effect of predators on prey *ma* ≤ 1), then the species with the highest biomass CV is always the perturbed species (Fig. 4C). Otherwise, if the perturbed species is not controlled by predation, then its prey is the species with the highest biomass CV (*e.g.*, species 1 when species 2 is perturbed for *ϵa* = 10 and *ma* = 10) as already observed by Barbier and Loreau (2019). In the case where species 1 disperses, species 2 is perturbed and species 1 is controlled by species 2 (food chain length equal to 2 or 4), species 1 has a higher biomass CV than species 2. Thus, the relative imbalance of biomass between the two patches is higher for species 1 than species 2, and species 1 dispersal depends on its biomass gradient. In the case where species 2 is controlled by species 3 (food chain length equal to 3) species 2 has a higher biomass CV, then the relative imbalance of biomass between the two patches is higher for species 2 than species 1, therefore, species 1 dispersal depends the biomass gradient of species 2 and an altered spatial synchrony is observed. Note that the case in which prey are perturbed and predators are able to disperse is completely symmetric (see Fig. S2-6 in the supporting information).

**Figure 4:**
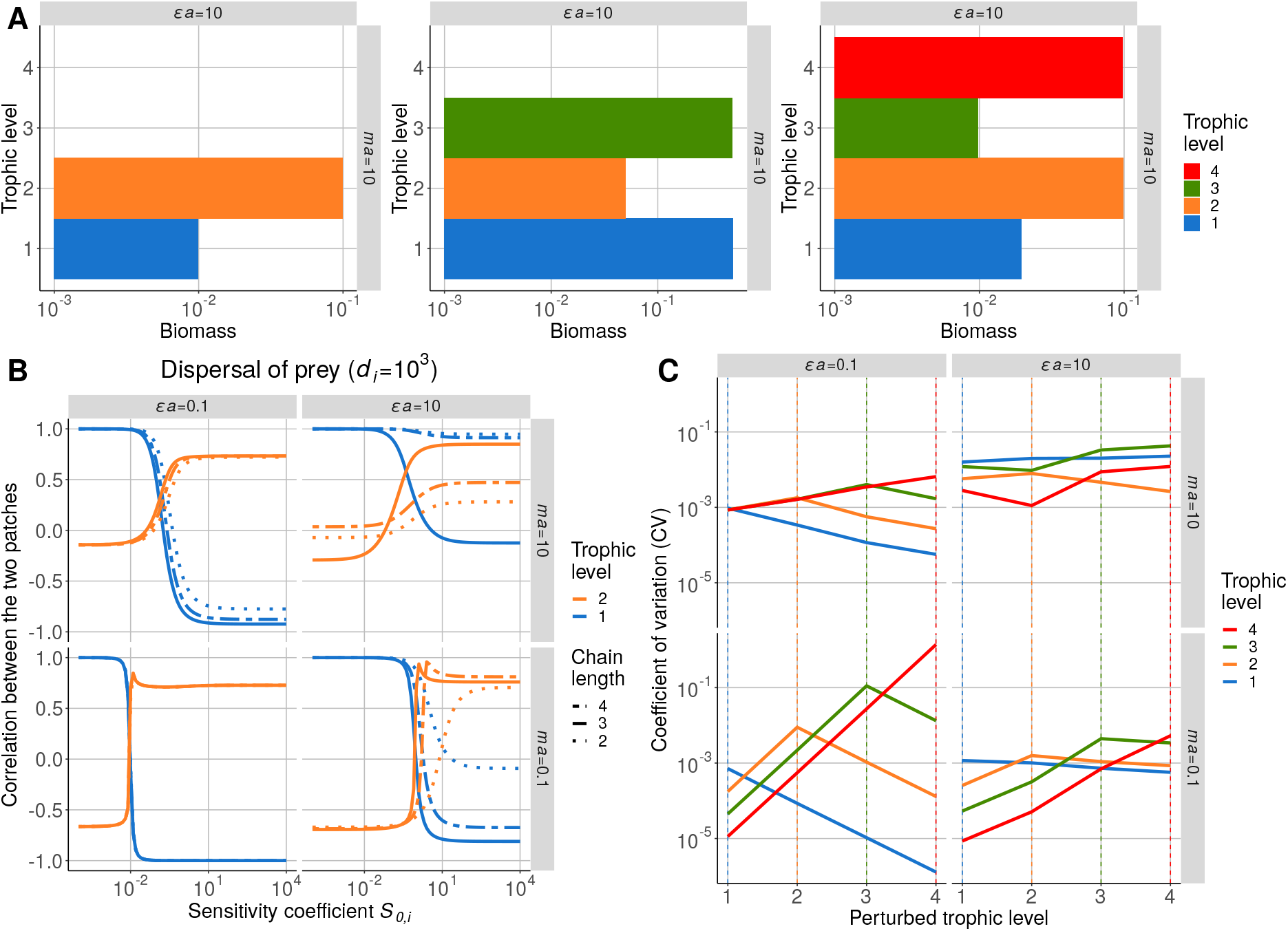
Effect of top-down control and food chain length on the influence of density-dependent dispersal. **A)** Biomass distribution when top-down control is strong (positive effect of prey on predators *ϵa* = 10 and negative effect of predators on prey *ma* = 10). **B)** Correlation between populations of trophic levels 1 and 2 in each patch when only species 1 disperses and its dispersal depends on the biomass of species 2. Correlation is calculated for increasing values of sensitivity coefficient *S*_0*,i*_ and for three different food chain lengths (*d_i_* = 10^3^). **C)** Biomass CV in a food chain with four species depending on which species is perturbed.

### Local demographic processes weight the different types of density-dependent dispersal

The main mechanism governing the effects of density-dependent dispersal on synchrony has been identified; thus, we can now study a more complex setup (Fig. 5). We consider a more realistic situation where all species are able to disperse at the same scaled rate *d_i_* and experience temporally correlated environmental perturbations in patch #1 (such as a drought or a bushfire, which similarly affect all species). In addition, we consider that dispersal components *f_self,i_*(*B_i_*), *f_pred,i_*(*B_i+1_*) and *f_prey,i_*(*B*_*i*−1_) are weighted as described by equations (7a-c). We detail the results for a positive effect of prey on predators *ϵa* = 0.1 and a negative effect of predators on prey *ma* = 10, although the response of the model for the entire parameter set is given by Fig. S2-14 in the supporting information. Note that the systematic effects of density-dependent dispersal and multiple independent perturbations are detailed in section S2-3 in the supporting information.

**Figure 5:**
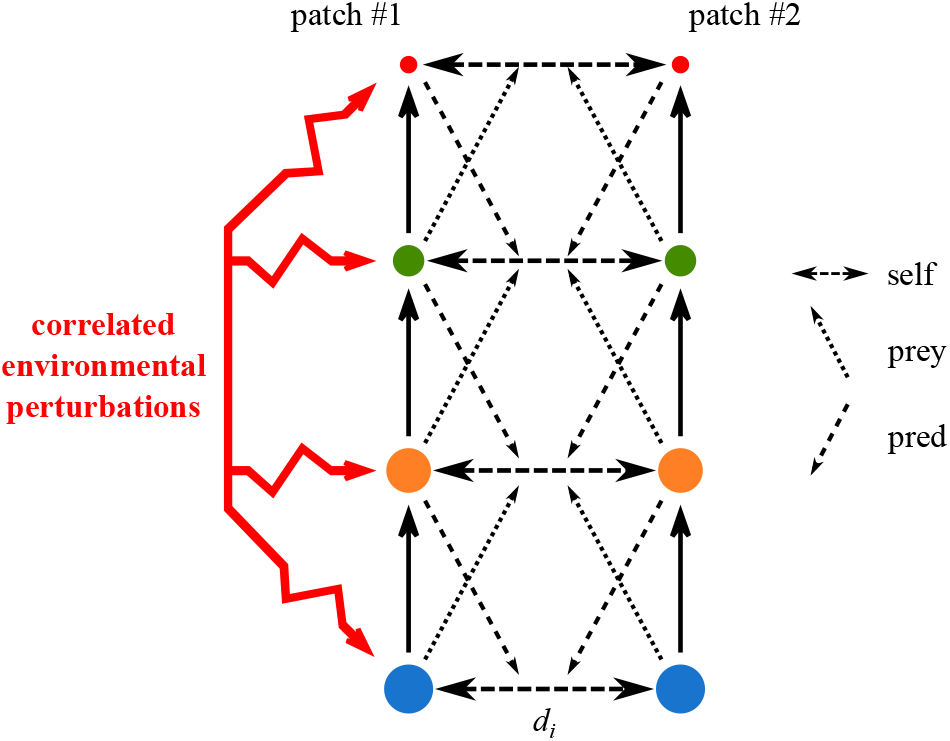
Schema of the realistic setup. Two patches sustain food chains with four trophic levels. All species in patch #1 experience perfectly correlated environmental perturbations (*z* = 1, see equation (9)) and all species disperse at the same scaled rate *d_i_*. The dispersal of each species depends on its own density (self), on the density of its prey (prey) and on the density of its predator (pred). Each dispersal component is weighted by the effects of self-regulation 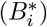, consumption 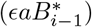 and predation 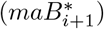 on the growth rate as detailed in equations (7a-c).

Again, the model with passive dispersal serves as a reference. The spatial synchrony in Fig. 6A is due to environmental perturbations, which affect more abundant species (primary producers), and dispersal, which has a stronger effect on the dynamics of less abundant species (see Box 3). In summary, basal species are relatively more perturbed, which leads to a bottom-up transmission of perturbations correlating all trophic levels in patch #1 (see Fig. S2-13D in the supporting information), and the top species mainly transmit perturbations to patch #2 because it is the less abundant (Fig. 7A and Fig. S2-13 in the supporting information). The underlying mechanisms have been detailed in Quévreux et al. (2021), and a description is provided in Fig. S2-13A in the supporting information. Therefore, we focus here on the mechanisms leading to the differences between Fig. 6A and 6B. Similar to the results presented above, density-dependent dispersal strongly alters spatial synchrony (Fig. 6B). At a low scaled dispersal rate *d_i_*, the top predator populations are correlated for passive and density-dependent dispersal. Thus, even though self and prey dependencies are balanced for top predators (Fig. 7B), prey dependency has an anti-correlating effect (Fig. 3A) while self-dependency has a correlating effect (Fig. S2-2A in the supporting information) and seems to have a stronger influence. These opposite effects of self-dependency and prey-dependency lead to a weaker transmission of perturbations compared to the case with passive dispersal, as showed by the lower biomass CV of top predators in patch #2 (Fig. 6C and 6D).

**Figure 6:**
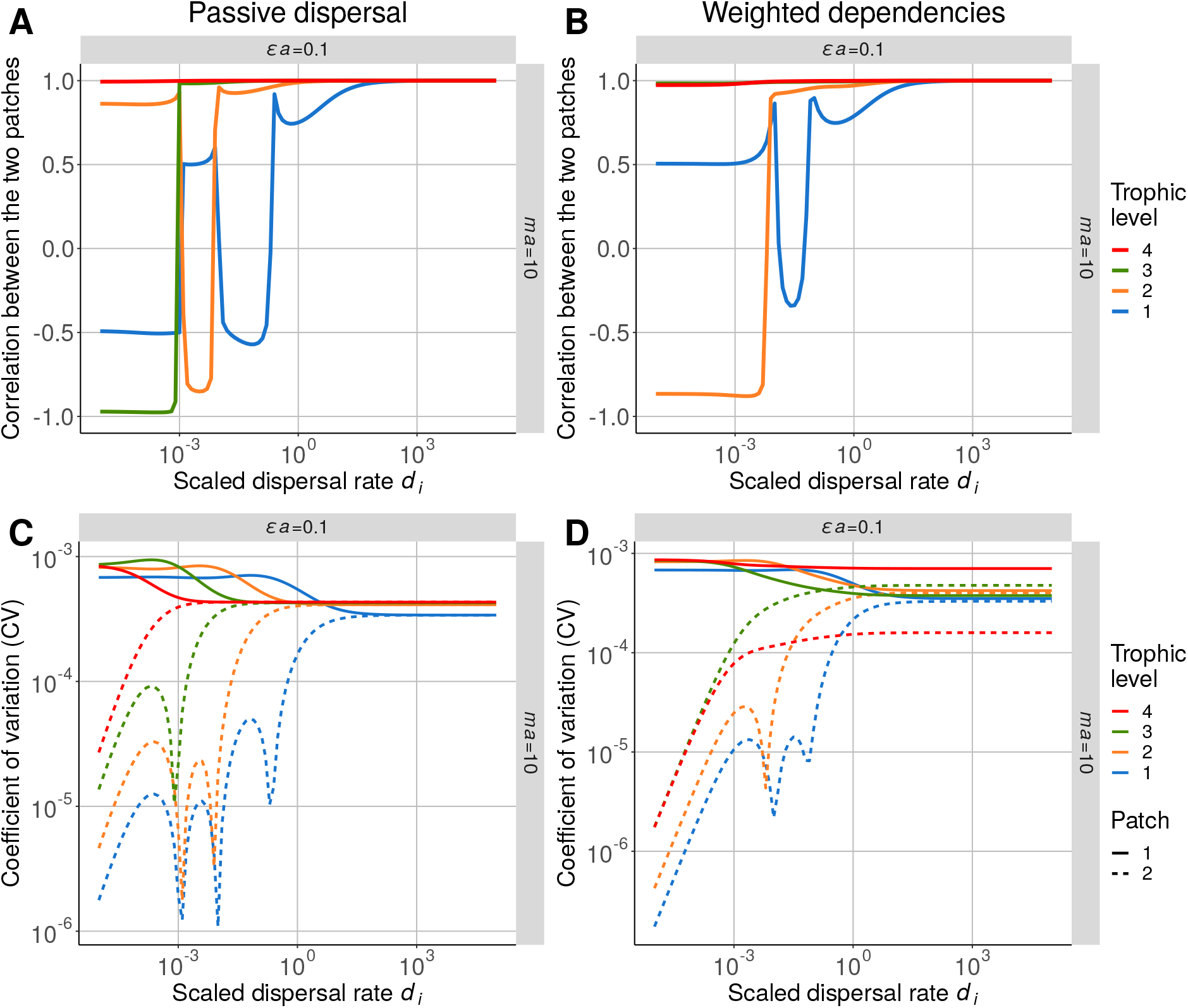
Response of a metacommunity where all species in patch #1 experience correlated environmental perturbations and are all able to disperse (positive effect of prey on predators *ϵa* = 0.1 and negative effect of predators on prey *ma* = 10). Correlation between populations and biomass CV in each patch when dispersal is **A) C)** passive or **B) D)** depends on self, prey and predator biomass (*S*_0*,self*_ = 10^3^, *S*_0*,prey*_ = 10^−3^ and *S*_0*,pred*_ = 10^3^) and is weighted by the associated demographic processes.

**Figure 7:**
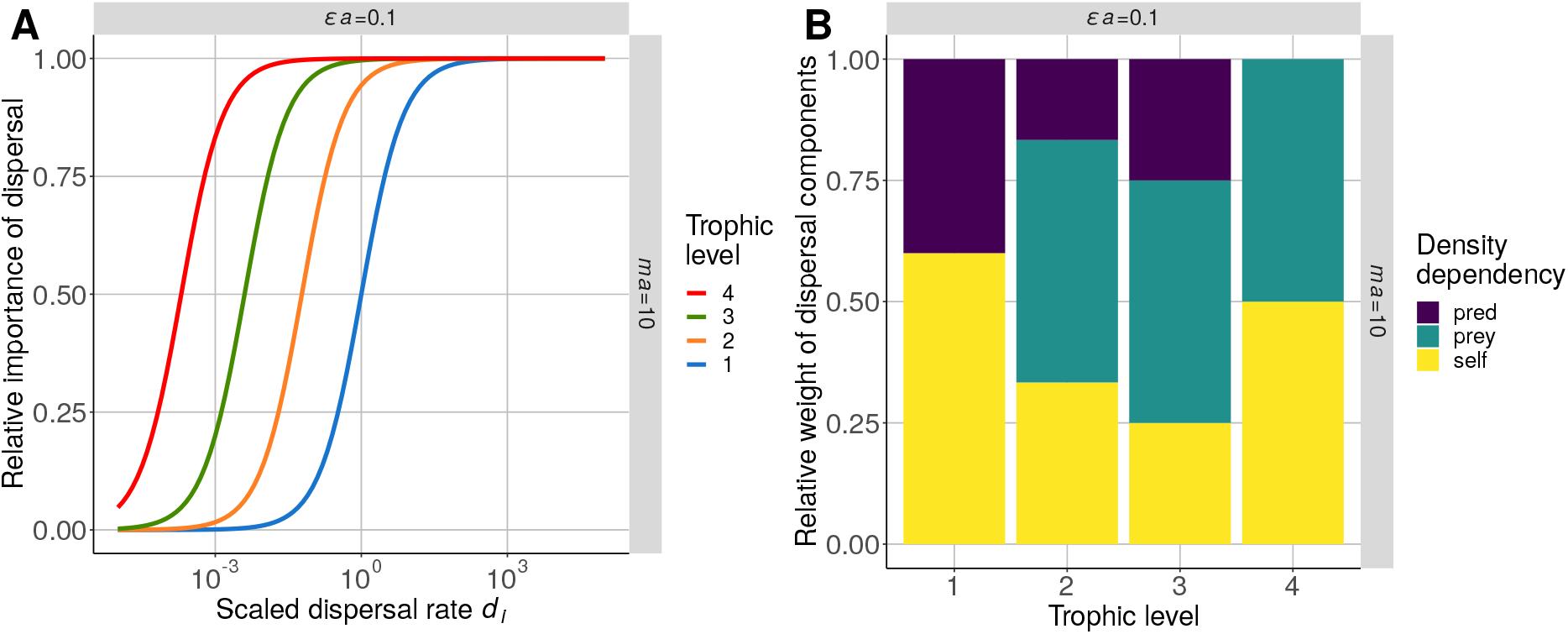
Importance of dispersal and its components in the overall dynamics for a positive effect of prey on predators *ϵa* = 0.1 and a negative effect of predators on prey *ma* = 10. **A)** Relative importance of dispersal processes compared to local demographic processes (see its definition in Box 3). **B)** Relative weight of each density dependency in the dispersal function (see equation (7)).

Although density-dependent dispersal does not alter the spatial synchrony of the top predators, we observe a complete change for carnivores (species 3) at low scaled dispersal rate *d_i_*. Indeed, carnivore populations are anticorrelated in the case of passive dispersal (Fig. 6A), while they are correlated in the case of density-dependent dispersal (Fig. 6B). Even if dispersal contributes less to the dynamics of carnivores than top predators (Fig. 7A), it strongly affects their spatial synchrony (see Fig. S2-18A and S2-18B in the supporting information). In fact, the biomass CV of top predators in patch #2 is equivalent to the biomass CV of carnivores (Fig. 6D) but higher in the case of passive dispersal (Fig. 6C). Thus, the top-down transmission in patch #2, which leads to the anticorrelation in Fig. 6A, is weaker and the correlating effect of dispersal influences relatively more the dynamics of carnivores in patch #2. In addition, the biomass CV of herbivores (species 2) is lower than the biomass CV of carnivores and top predators. Thus, even if prey-dependency weights for 50% in the dispersal function, self and predator dependencies, which have a correlating effect in this setup, have a stronger influence. Self density-dependent dispersal correlates populations in the same way as passive dispersal because the vertical transmission of perturbations cannot be bypassed (Fig. 1B and Fig. S2-8 in the supporting information). Here, predator density-dependent dispersal is correlating because perturbations of top predators and carnivores are temporally correlated. For instance, an increase in biomass of the two species in patch #1 leads to an increase in carnivore biomass in patch #2 because carnivores avoid predation by emigrating (see the time dynamics in Fig. S2-15 in the supporting information).

For higher dispersal rates, the spatial synchrony of lower trophic levels is identical to the case with passive dispersal because when dispersal starts being important for herbivores and primary producers (Fig. 7A), their biomass CV in patch #1 is higher than the biomass CV of their predators or their prey. As demonstrated by Fig. 4, dispersal follows the biomass gradient of the dispersing species and is equivalent to passive dispersal.

## Discussion

Most studies consider passive dispersal for the sake of simplicity, although our results show that density-dependent dispersal can reverse the spatial synchrony compared to the one predicted by these studies. We find that when dispersal depends on the biomass of several species (*e.g.*, predators or prey), the species with the highest biomass CV drives the dispersal of the dispersing species. This means that dispersal is determined by the biomass imbalance of predators or prey between the two patches instead of the biomass imbalance of the dispersing species. However, this rule can be modulated by the trade-off between foraging for food and predator avoidance since species disperse in response to these biological needs to maximise their fitness. Our results are overall consistent with Li et al. (2005) and Hauzy et al. (2010), although the existing discrepancies are due to the presence of limit cycles in their systems and the observed equilibrium reached by our system, as explained below.

### Density-dependent versus passive dispersal

We are able to disentangle all the mechanisms acting in our system to explain the spatial synchrony in metacommunities. For instance, in the metacommunity presented in Fig. 3A, the dispersal of predators depends on their own biomass and on prey biomass (see equation equation (4b)). On the one hand, passive dispersal tends to correlate populations, as reported by previous studies considering passive dispersal (Bjørnstad et al., 1999; Lloyd and May, 1999). On the other hand, the effect of prey density anticorrelates predator populations because it amplifies the bottom-up effect of prey. Our results show that the species with the highest biomass CV determines which of these two effects drives dispersal; thus, we can easily predict the effect of density-dependent dispersal based on the distribution of biomass CV.

The consequences of density-dependent dispersal on stability are different in our study compared to previous studies. Hauzy et al. (2010) found that density-dependent and passive dispersal can increase or decrease regional biomass CVs, while they have no effect in our model (Fig. S2-16 in the supporting information). This discrepancy should be due to the different regimes reached by each study: limit cycles for Hauzy et al. (2010) and equilibrium in our case. As dispersal has no cost in these models (*i.e.*, no additional mortality), the emigration from a patch is equal to the immigration to the other patches, making the dynamics of the total biomass independent of dispersal (Wang et al., 2015). Despite the absence of effects of dispersal on the stability of the total biomass of each species in our model, density-dependant dispersal alters biomass CVs at the local scale with an increase or a decrease depending on the considered trophic level and the weight attributed to each dependency (Fig. S2-14C and S2-14C and Fig. S2-17 in the supporting information). However, the potential stabilisation (*i.e.*, reduction in biomass CV) in one patch is balanced by destabilisation in the other patch because dispersal does not affect the stability of the total biomass of each species.

The effects of density-dependent dispersal on metacommunity dynamics cannot be easily assessed empirically because the effects of local trophic dynamics cannot be disentangled from the effects of density-dependent dispersal. Various levels of predator cues can be tested on prey metapopulations to trigger density-dependent dispersal (Hauzy et al., 2007; Fronhofer et al., 2018; Harman et al., 2020), although such a treatment removes intrapatch predator-prey dynamics. Conversely, the direct addition of predators necessarily triggers density-dependent dispersal (Hauzy et al., 2007; Vasseur and Fox, 2009) and does not allow for the study of a reference trophic metacommunity with passive dispersal. Therefore, we recommend considering density-dependent dispersal in metacommunity models because it is indissociable from trophic interactions.

### Dispersal and top-down and bottom-up control

The central mechanism governing the spatial synchrony in our metacommunity model is the distribution of biomass CVs among species. Indeed, density-dependent dispersal is driven by the species with the highest biomass CV (Fig. 8A). For instance, in a system with three species in which only herbivores disperse, predator density-dependent dispersal prevails only if predators have the highest biomass CV (Fig. S2-12 in the supporting information). We can identify two mechanisms that control the species that has the highest biomass CV. First, a particular species can be more affected by perturbations because it is targeted (*e.g.* harvesting) or more sensitive to perturbations. Arnoldi et al. (2019) showed that abundant species are more affected by environmental perturbations and Barbier and Loreau (2019) showed that a few ecological and physiological parameters determine biomass distribution among species in food chains (Fig. S2-13A in the supporting information). Thus, the sensitivity of species to perturbations can be efficiently forecasted.

**Figure 8:**
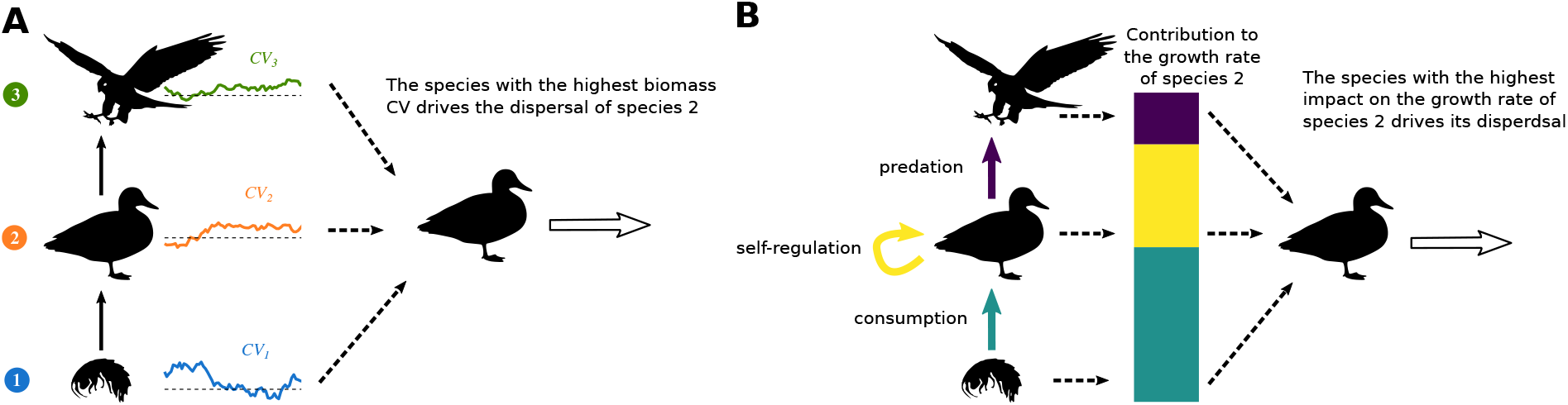
Mechanisms determining which type of density-dependent dispersal drives the dispersal of species 2 in a three trophic levels food chain. **A)** The species with the highest biomass CV drives the dispersal of species 2. For instance, if predators (species 3) have the highest biomass CV, predator density-dependent dispersal is dominant and species 2 disperse depending on the abundance of predators in the two patches. **B)** The species that has the highest impact on the growth rate of species 2 drives its dispersal. For instance, if species 2 has a low mortality due to predators, predator density-dependent dispersal has a minor impact on species 2 dispersal. The final outcome of dispersal is a balance between these two mechanisms. For instance, even if predators have a high dispersal rate, they will not necessarily have a major impact on the dispersal of species 2 if they have a low impact on its growth rate.

Second, biomass CV distribution strongly depends on food chain length in the case of tropic cascades, which occur when interspecific interactions are relatively stronger than intraspecific interactions (Fig. 4). Shanafelt and Loreau (2018) showed that the biomass CV of controlled species is high because their biomass is much lower than the biomass of other species. In the example presented in Figs. 3 and 4, which includes two species where only prey are able to disperse, prey are controlled and have the highest biomass CV. Thus, prey dispersal does not depend on predator abundance, while predators have a strong impact on prey mortality. This paradox is restricted to systems with a stable equilibrium because Hauzy et al. (2010) found an effect of predators on prey dispersal in a similar system that displayed limit cycles. A comparison between stable equilibrium and limit cycles is discussed in the following section.

Although identifying the species that has the highest biomass CV is key to understanding spatial synchrony from a mathematical point of view, the contribution of each type of density-dependency to the dispersal function can be modulated by biological factors (French and Travis, 2001). Here, we propose that each dependency should be weighted by the relative contribution of the associated local demographic process to the overall growth rate (Fig. 8B). In fact, prey live in a landscape of fear (Laundré et al., 2010) in which they have to balance foraging for food with predator avoidance (Gilliam and Fraser, 1987; Preisser et al., 2005; Preisser and Bolnick, 2008) with a potential cascading effect in food chains (Bestion et al., 2015; Blubaugh et al., 2017). Such adaptive responses of species can be compared to adaptive foraging where consumers adapt their feeding effort among prey to maximise their resource intake (Loeuille, 2010), thus promoting species coexistence and food web stability by avoiding overexploitation (Kondoh, 2003; Uchida and Drossel, 2007; Heckmann et al., 2012). In our model, this optimisation of dispersal does not stabilise the entire population because dispersal does not affect biomass dynamics at the metacommunity scale (Fig. S2-17 in the supporting information). However, it potentially alters spatial synchrony because the distribution of biomass CV among species does not necessarily fit the relative importance of demographic processes (Fig. S2-12 in the supporting information). Thus, considering both mechanisms underlying perturbation transmission and ecological factors triggering dispersal is required to properly predict the response of metacommunities to perturbations.

### To equilibrium and beyond

The major difference between our work and previous studies is that we consider stable equilibria instead of limit cycles (Li et al., 2005; Hauzy et al., 2010; Abrams and Ruokolainen, 2011; Fronhofer et al., 2018), which implies that different mechanisms control population synchrony and community stability.

First, the nonlinearity of density-dependent dispersal functions is key in systems where species biomasses experience large variations because it leads to substantial changes in dispersal rates over time. In turn, these strong changes deeply alter the overall dynamics of the metacommunity. In our system in the vicinity of equilibrium, these functions are equivalent to linear functions since the system is near equilibrium and affected by small perturbations (Fig. S2-19 in the supporting information). Thus, the variations in biomass of various species do not lead to dramatic variations in dispersal rates but simply add new pathways for perturbation transmission. As explained earlier, if the biomass CV of prey or predators is higher than the biomass CV of the dispersing species, perturbations are transmitted between patches by density-dependent effects and bypass the classic vertical transmission, which relies on trophic interactions. Therefore, the distribution of biomass CVs is the main mechanism governing synchrony in metacommunities near equilibrium.

Second, phase locking is one of the main mechanisms governing the dynamics of systems displaying limit cycles and coupled oscillators in general (Jansen, 1999; Lloyd and May, 1999; Goldwyn and Hastings, 2008; Vasseur and Fox, 2009). In meta-communities, each patch is an oscillator coupled to the other patch through dispersal. High dispersal tends to synchronise the phases of oscillators, while low dispersal Jansen (1995) and Hauzy et al. (2010), asymmetry between patches (McCann et al., 1998; Vasseur and Fox, 2007; Goldwyn and Hastings, 2009) or connectivity of several patches (Marleau et al., 2014; Hayes and Anderson, 2018; Guichard et al., 2019) can lead to asynchrony. In the case of synchronised patches, density-dependent dispersal decouples the various communities and promotes asynchrony (Li et al., 2005; Hauzy et al., 2010). Phase locking leads to synchrony patterns completely different from our results. For instance, a higher dispersal of prey than predators in Hauzy et al. (2010) leads to synchrony of both predators and prey, while our results show that it can lead to the asynchrony of predators if they are perturbed (Figs. 3 and S2-1 in the supporting information). In fact, perturbations affecting systems at equilibrium lead to responses consistent with the trophic cascade framework (Wollrab et al., 2012; Quévreux et al., 2021). These discrepancies demonstrate that synchrony in trophic metacommunities is highly sensitive to the type of dispersal, the state of the system (equilibrium, limit cycles, chaos) and the presence of perturbations.

Taken together, our results and these studies outline two facets of synchrony in metacommunities because biomass variability can be generated internally by predator-prey oscillators or externally by perturbations. The combined effect of limit cycles and stochastic perturbations can lead to counterintuitive results. For instance, Vasseur and Fox, 2007 showed that spatially correlated environmental perturbations decrease the synchrony between the two consumers in a diamond-shaped food web but also decrease the temporal variability of their biomass dynamics. Finally, in between small perturbations and limit cycles lie large perturbations whose effects have been poorly explored. Indeed, large perturbations are generated externally, although they push the system too far from equilibrium for the linear approximation to hold. Thus, if the system is at equilibrium but close to the Hopf bifurcation, large perturbations will lead to dampened oscillations (Rooney et al., 2006) with characteristics close to that of limit cycles. Considering all these types of variability and perturbations should help us to determine the entire spectrum of dynamics of trophic metacommunities and to better grasp their response to anthropogenic perturbations.

## Conclusions and perspectives

Dispersal is a complex process that is much more than a simple balancing of biomass between distant populations. By considering density-dependent dispersal, we showed that the cross effects of predators and prey on their dispersal can deeply shape synchrony in metacommunities in response to perturbations. In our model, the different types of density-dependent dispersal have additive effects as demonstrated experimentally by Fronhofer et al. (2018); however, future models could consider multiplicative effects as well. For instance, French and Travis (2001) reported that parasitoid wasps disperse more when the parasitoid:host ratio is high because it reflects an increase in intraspecific competition when resources are scarce. We hope our results will help future studies better describe the complexity of processes involved in dispersal and thus improve our understanding of metacommunity dynamics.

## Data accessibility

The R code of the simulations and the figures is available on Zenodo (doi:10.5281/zenodo.5013867).

## Acknowledgement

This work was supported by the TULIP Laboratory of Excellence (ANR-10-LABX-41) and by the BIOSTASES Advanced Grant, funded by the European Research Council under the European Union’s Horizon 2020 research and innovation programme (666971). We also thank Matthieu Barbier, Julien Cote and Delphine Legrand for their useful discussions and comments, and the two anonymous reviewers who greatly helped us to improve the readability and the clarity of this manuscript.

## S1 Complementary material and methods

### S1-1 Parameters

**Table S1-1:** Distribution of *m* and *a* leading to the desired values of *ma* and *ϵa*.

| | $\epsilon a = 0.1$ | $\epsilon a = 1$ | $\epsilon a = 10$ |
| --- | --- | --- | --- |
| $ma = 10$ | $m = 65$<br>$a = 0.15$ | $m = 6.5$<br>$a = 1.5$ | $m = 0.65$<br>$a = 15$ |
| $ma = 1$ | $m = 6.5$<br>$a = 0.15$ | $m = 0.65$<br>$a = 1.5$ | $m = 0.065$<br>$a = 15$ |
| $ma = 0.1$ | $m = 0.65$<br>$a = 0.15$ | $m = 0.065$<br>$a = 1.5$ | $m = 0.0065$<br>$a = 15$ |

### S1-2 Demographic and environmental perturbations

This section summarises the demonstration of the scaling of demographic and environmental perturbations with species abundance (see Lande et al., 2003 for more details).

The growth of a population depends on the fitness *w_i_* of each individual *i*. This fitness can be decomposed into the expected fitness *µ_w_* of the species depending on environmental conditions and the individual deviation *δ_i_*. The expected value of *w_i_* is equal to *µ_w_* (*E*(*w_i_*) = *µ_w_* and *E*(*δ_i_*) = 0).

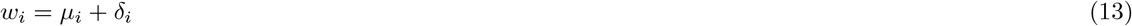

Then, the growth rate *λ* of the population, which contains *N* individuals, is the mean of the *w_i_*.

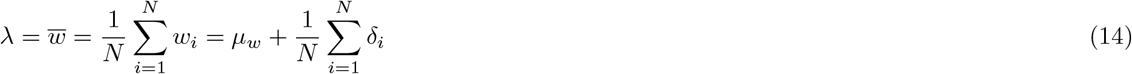

*µ_w_* and *δ_i_* are independent random variables whose variance are respectively 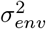 and 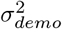. Thus, we can calculate the variance of *λ*.

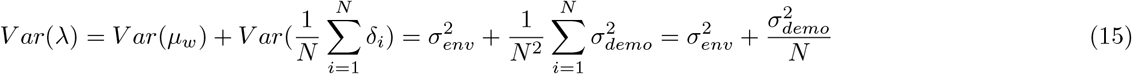

As the growth of the population is defined by *λN*, we get

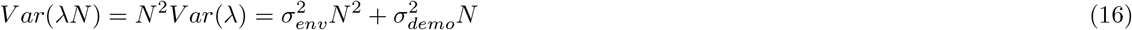

Thus, the variance of the population due to a synchronised response of all the individuals is equal to 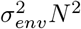 while the variance of the demographic noise is equal to 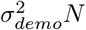. In conclusion, we can represent the demographic perturbation by *σ_demo_B*^0.5^*dW* and the environmental perturbation by *σ_env_BdW* as in equation (9) in the main text.

### S1-3 Biomass at equilibrium

The system of equations (4a) and (4b) at equilibrium cannot be solved analytically. Therefore, we calculate analytically the equilibrium biomass of the system without dispersal by solving the following equations:

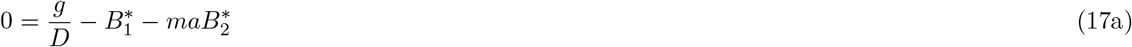

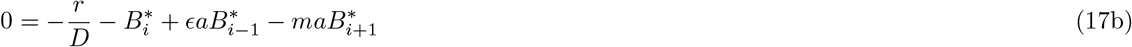

This can be expressed as a matrix product:

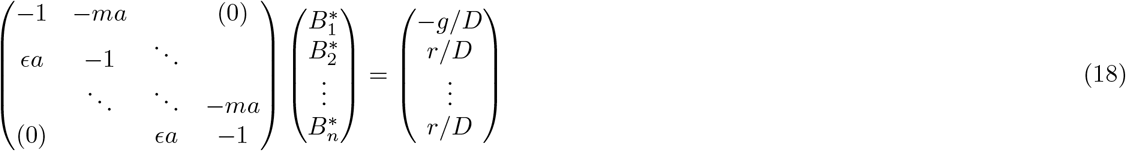

solved by the tridiagonal solver algorithm of the GNU Scientific Library version 2.5 (Galassi, 2009). Then, these values initialise the multidimensional root-finder algorithm of the GNU Scientific Library version 2.5 (Galassi, 2009) that solve the following system to find the equilibrium biomasses of the metacommunity.

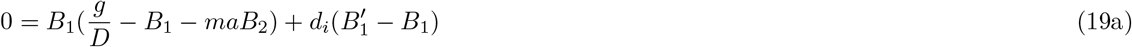

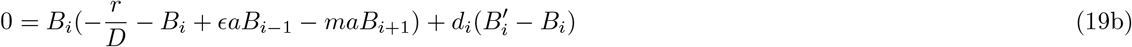

These equations have the same solutions that equations (4a) and (4b) but the absence of *m*_*i*−1_, which can be very low or high depending on the value of *m*, greatly increases the precision of the algorithm.

### S1-4 Jacobian matrix

The general system with *S* species is defined as follows:

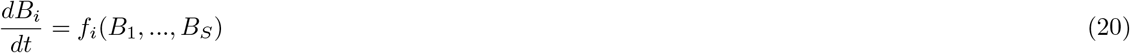

*B^∗^* defines the equilibrium at which the community matrix (or Jacobian matrix) *J* is defined as follows:

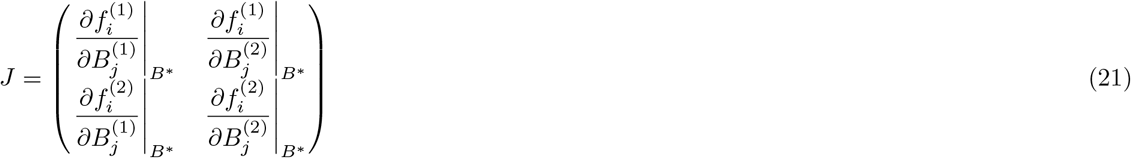

where 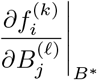 represents the effect of species from patch *£* on dynamics of species from patch *k*. For simplicity, *J* can be split into blocks, such as the following:

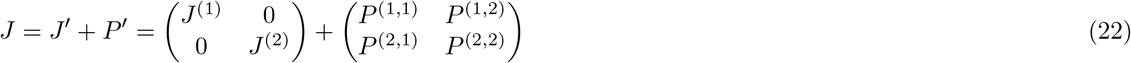

where *J* ^(*k*)^ is the Jacobian matrix of community *k* without dispersal and *P* ^(*k,*)^ is the sub-dispersal matrix (with size *S* × *S*). *J* ^(*k*)^ is defined as follows:

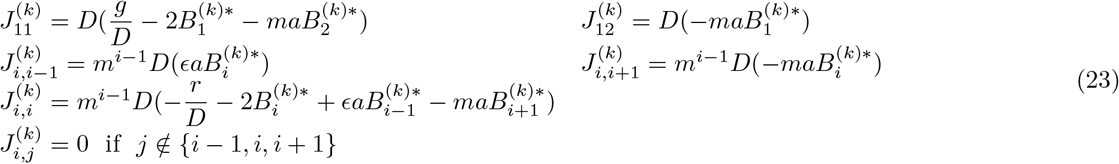

If *k* = *ℓ*

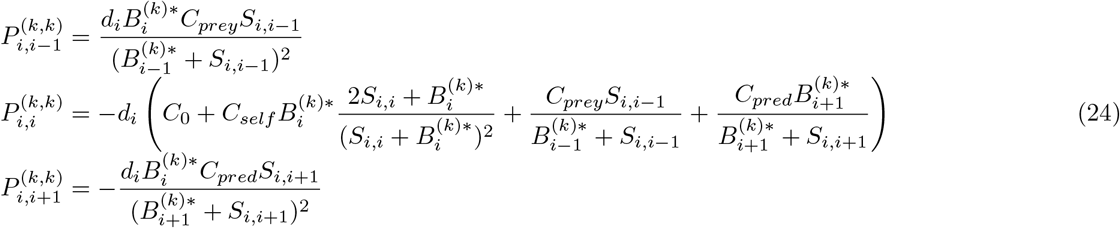

If *k* I= *ℓ*

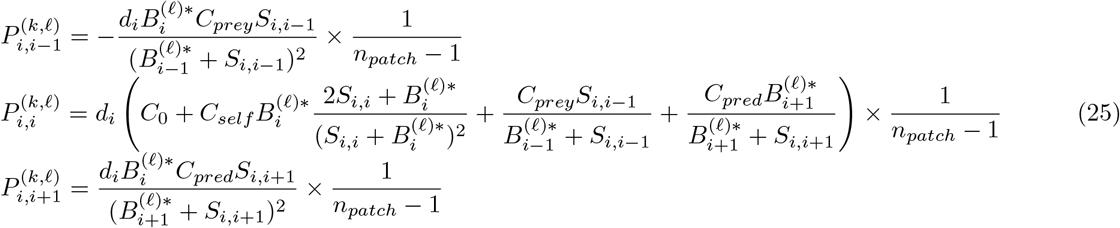

### S1-5 Linearisation of the system and resolution of the Lyapunov equation

#### S1-5-1 Linearisation of the system

The system of equations (4a) and (4b) can be linearised in the vicinity of *B∗*:

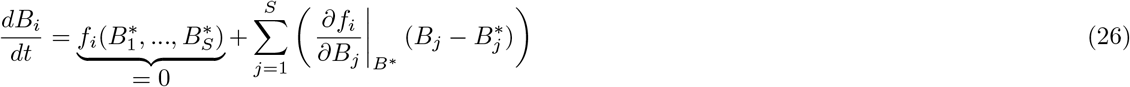

Thus, by setting 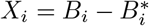 the deviation from equilibrium, we have:

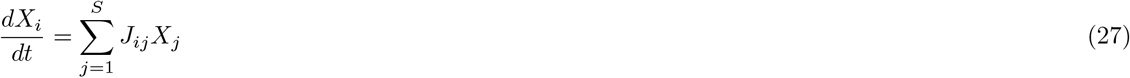

Then, we can consider small perturbations defined by 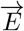 whose effects on 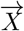 are defined by the matrix *T* (Arnoldi et al., 2016). We get the linearised version of equation (9):

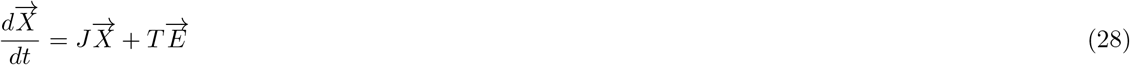

The elements of 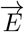 are defined by stochastic perturbations *E_i_* = *σ_i_dW_i_* with *σ_i_* their standard deviation and *dW_i_* a white noise term with mean 0 and variance 1. In this study, each species can receive three types of perturbation scaling with each biomass 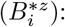 : exogenous if *z* = 0, demographic if *z* = 0.5 and environmental if *z* = 1 (see section S1-2). Thus, 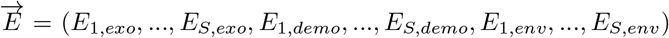 as it contains the white noise term of each type of perturbation for each species. *T* contains three blocks of diagonal matrices of size *S* corresponding to each type of perturbation.

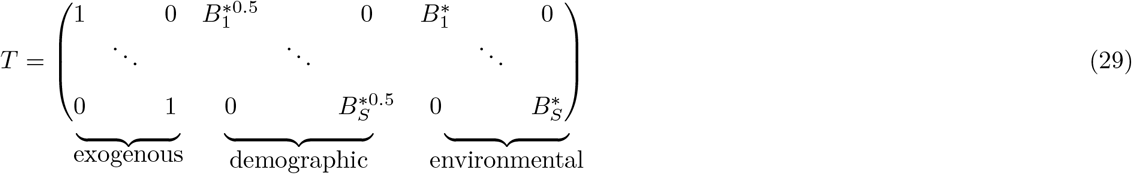

Thus, the matrix product 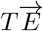 results in the product of the white noise and the biomass scaling as in equations (4a) and (4b) in the main text. Moreover, each species can receive simultaneously one perturbation of each type.

#### S1-5-2 Resolution of the Lyapunov equation

In the vicinity of equilibrium, the Lyapunov equation links the variance-covariance matrix *VE* of the perturbation vector 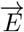 to the variance-covariance matrix *C^∗^* of species biomasses (see the appendix of Wang et al. (2015) for more details on the Lyaponov equation).

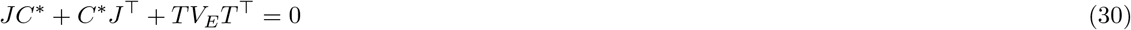

The diagonal elements of *V_E_* are equal to 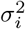 (variance of the white noises) and the non-diagonal elements are equal to zero if perturbations are independent (what we consider in this study). ⊤ is the transpose operator.

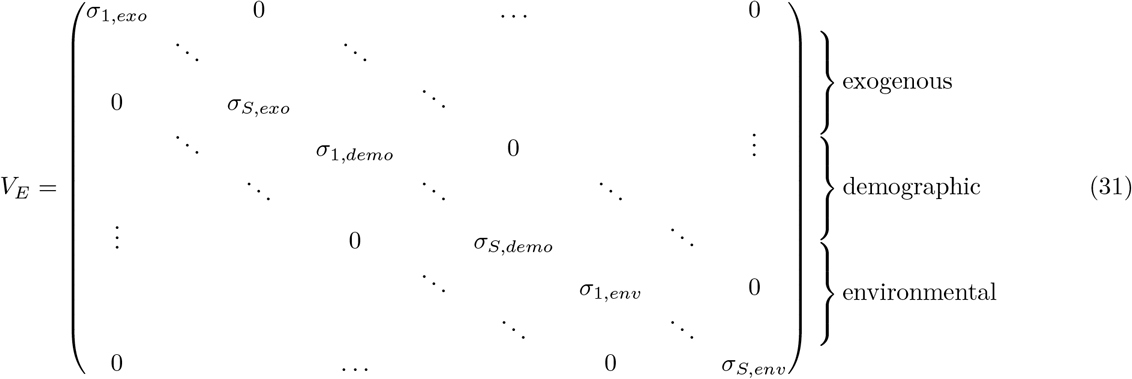

*C*^∗^ can be calculated using a Kronecker product (Nip et al., 2013). The Kronecker product of an *m* × *n* matrix *A* and a *p* × *q* matrix *B* denoted *A* ⊗ *B* is the *mp* × *nq* block matrix given by:

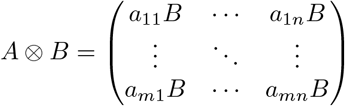

We define 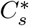 and (*TV_E_T^⊤^*)_*s*_ the vectors stacking the columns of *C^∗^* and *TV_E_T^⊤^* respectively. Thus, equation (30) can be rewrite as:

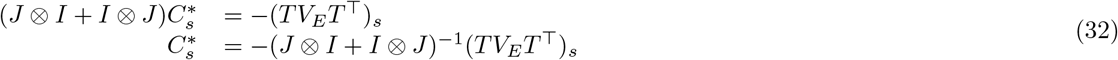

### S1-6 Numerical resolution of stochastic differential equations

The same results can be obtained by simulating the temporal dynamics of the system. Equation (9) in the main text can be solved by using the Euler-Maruyama method by computing the discretised equation:

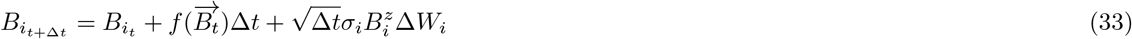

With ∆*t* the time step and ∆*W_i_* a displacement drawn from a Gaussian distribution with 0 mean and variance 1.

## S2 Complementary results

### S2-1 Dispersal depending on focal species abundance

**Figure S2-1:**
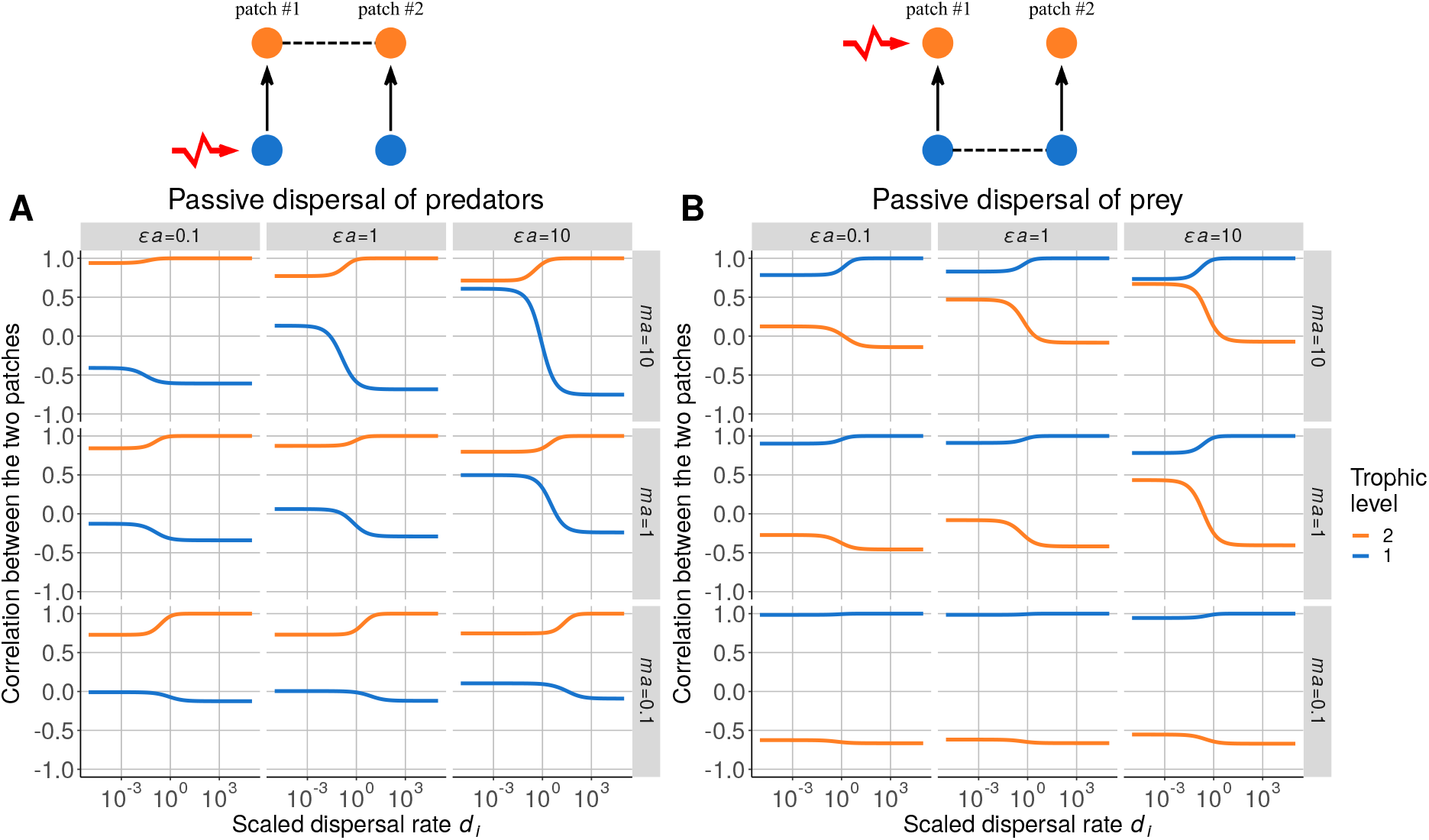
Correlation between populations in each patch in the case of passive dispersal of a single species (classic mass effect). **A)** Prey are perturbed, and predators are able to disperse. **B)** Predators are perturbed, and prey are able to disperse.

The setup described in Fig. S2-1 corresponds to the model developed in Quévreux et al. (2021) and summarised in Box 1 in the main text. The populations of the dispersing species are perfectly correlated at high scaled dispersal rate *d_i_*, while the nondispersing species populations are anticorrelated.

As the perturbed species does not disperse, perturbations have a bottom-up transmission in one patch and a top-down transmission in the other patch (Fig. S2-1A). This dynamics leads to different correlation patterns between species within each patch, which ultimately determine whether the two populations of the same species are correlated or anticorrelated (Fig. S2-1B and S2-1C, see also Quévreux et al. (2021) who thoroughly describe the mechanisms).

Dispersal dependency on the biomass of the dispersing species (self-dependency) leads to the same results as passive dispersal (Fig. S2-1 and Fig. S2-2A and S2-2B). Indeed, varying the sensitivity coefficients *S*_0*,i*_ does not change the spatial synchrony (Fig. S2-2C and S2-2D) compared to the low values of *S*_0*,self*_ which is equivalent to the case with passive dispersal (*f_self,i_*(*B_i_*) ≃ 1, see equation (6a)).

### S2-2 Dispersal depending on prey or predator abundance

**Figure S2-2:**
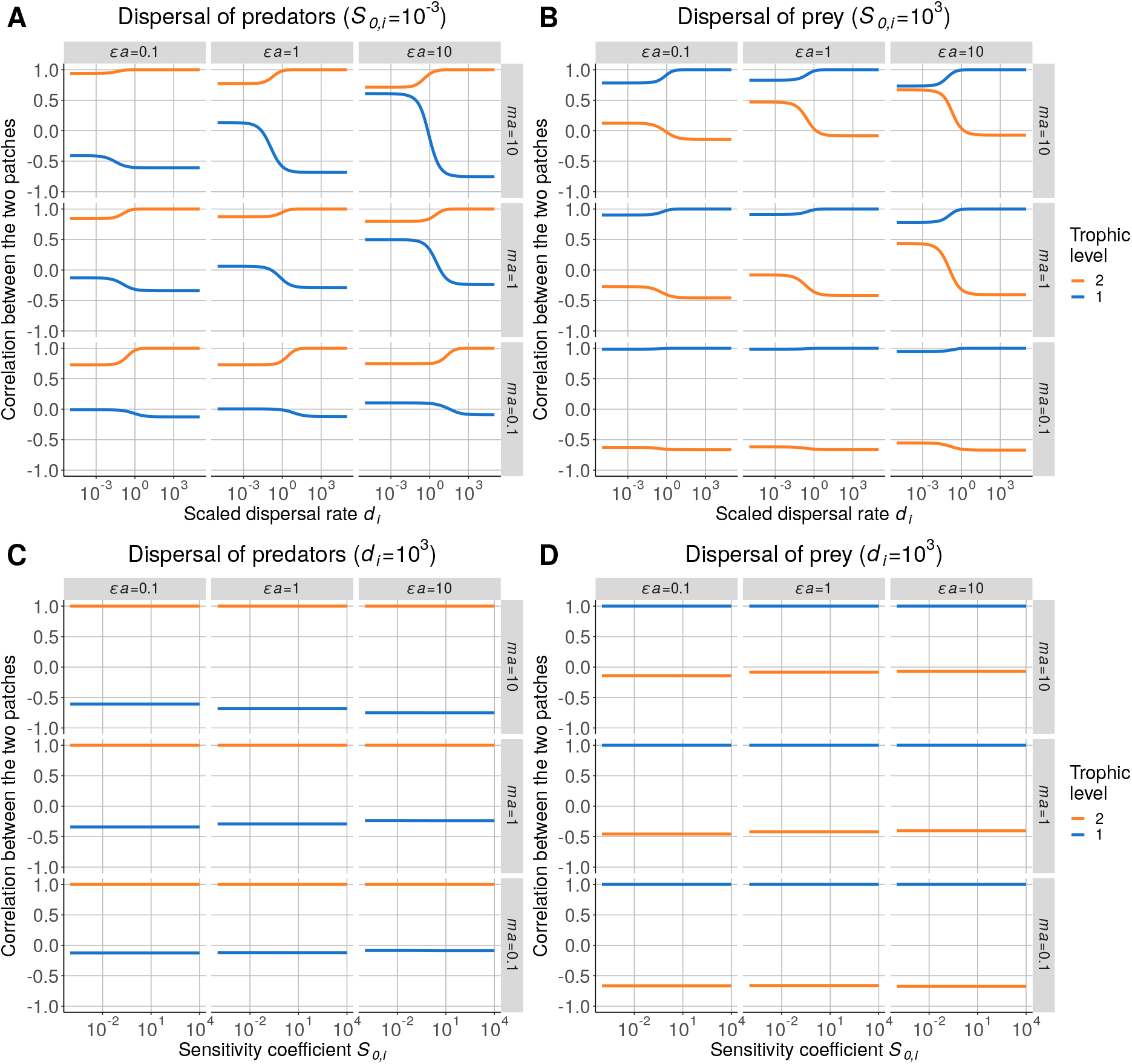
Correlation between populations in each patch when dispersal depends on the biomass of the dispersing species for increasing values of scaled dispersal rate *d_i_*. **A)** Perturbed prey and dispersing predators (*S*_0_ = 10^−3^). **B)** Perturbed predators and dispersing prey (*S*_0_ = 10^3^). Spatial synchrony for increasing values of biomass sensitivity *S*_0_. **C)** Perturbed prey and dispersing predators (*d_i_* = 10^3^). **D)** Perturbed predators and dispersing prey (*d_i_* = 10^3^).

In this setup, predator dispersal depends on prey biomass density, primary producers are perturbed in patch #1 and only predators are able to disperse. Thus, equation (5) only depends on *f*_*prey*,2_(*B*_1_) (*C*_0*,i*_ = 0, *C_self,i_* = 0 and *C_pred,i_* = 0). In this case, the spatial synchrony is inverted compared to the case with passive dispersal for a sensitivity coefficient *S*_0*,i*_ = 10^−3^ (Fig. S2-1A and S2-3A) while it is similar for a sensitivity coefficient *S*_0*,i*_ = 10^3^ (Fig. S2-3C). In fact, *f_prey,i_*(*B*_*i*−1_) ∼ 1 for a sensitivity coefficient *S*_0*,i*_ = 10^3^, which leads to a dispersal function *F_i_*(*B_i−1_*, *B_i_*, *B*_*i*+1_) similar to passive dispersal, while *f_i,prey_* ∼ *S*_0*,i*_/B_i−1_ for a sensitivity coefficient *S*_0*,i*_ = 10^−3^ (equation (6c) and Fig. S2-4).

**Figure S2-3:**
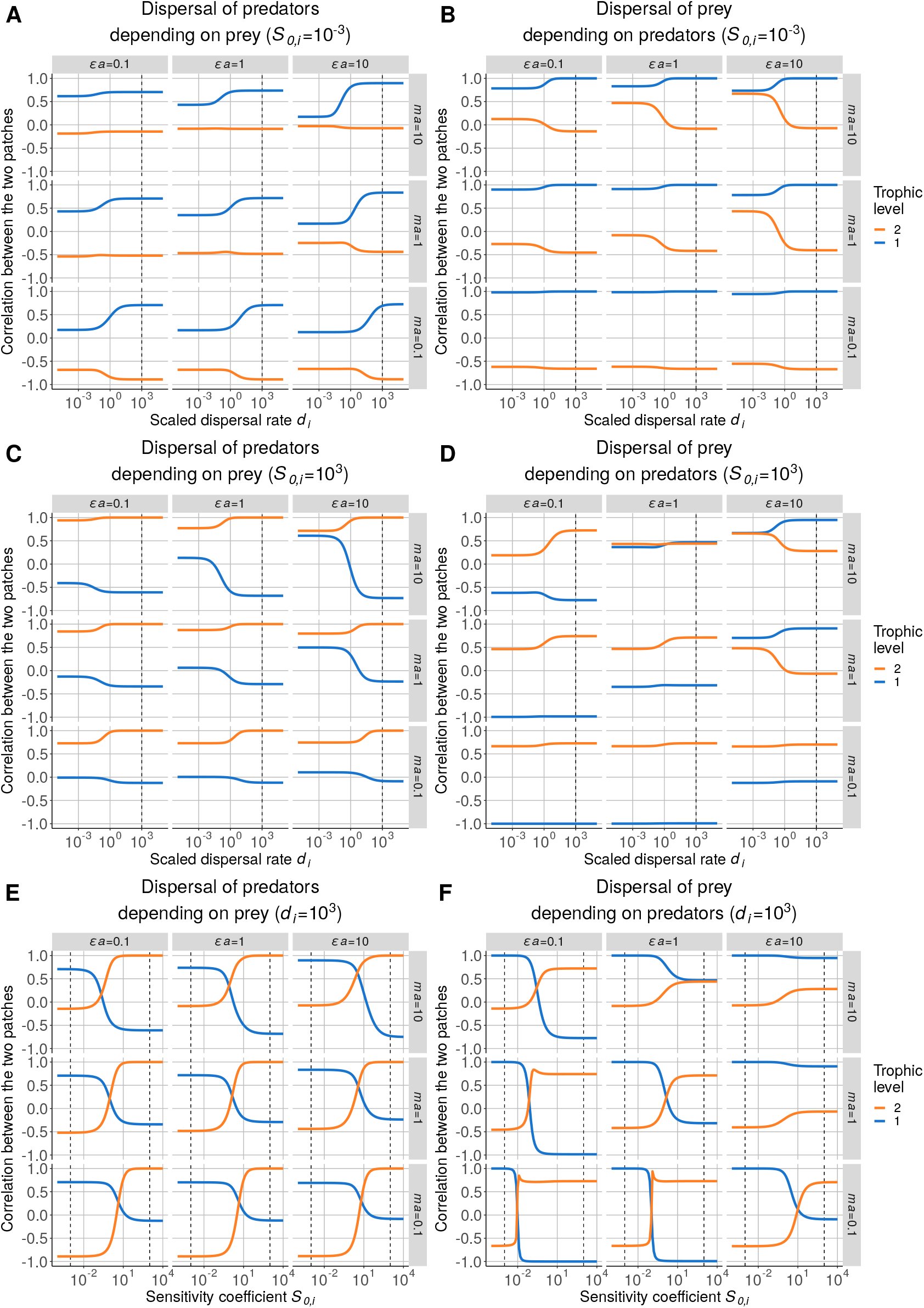
Correlation between populations in each patch when predator dispersal depends on prey biomass: **A)** when *S*_0*,i*_ = 10^−3^ and **B)** when *S*_0*,i*_ = 10^3^ for increasing values of scaled dispersal rate *d_i_* and **E)** increasing values of sensitivity coefficient *S*_0*,i*_ (*d_i_* = 10^3^). Correlation between populations in each patch when prey dispersal depends on predator biomass: **B)** when *S*_0*,i*_ = 10^−3^ and **D)** when *S*_0*,i*_ = 10^3^ for increasing values of scaled dispersal rate *d_i_* and **F)** increasing values of sensitivity coefficient *S*_0*,i*_ (*d_i_* = 10^3^). The dashed lines represent the values of *d_i_* and *S*_0*,i*_ between graphs.

**Figure S2-4:**
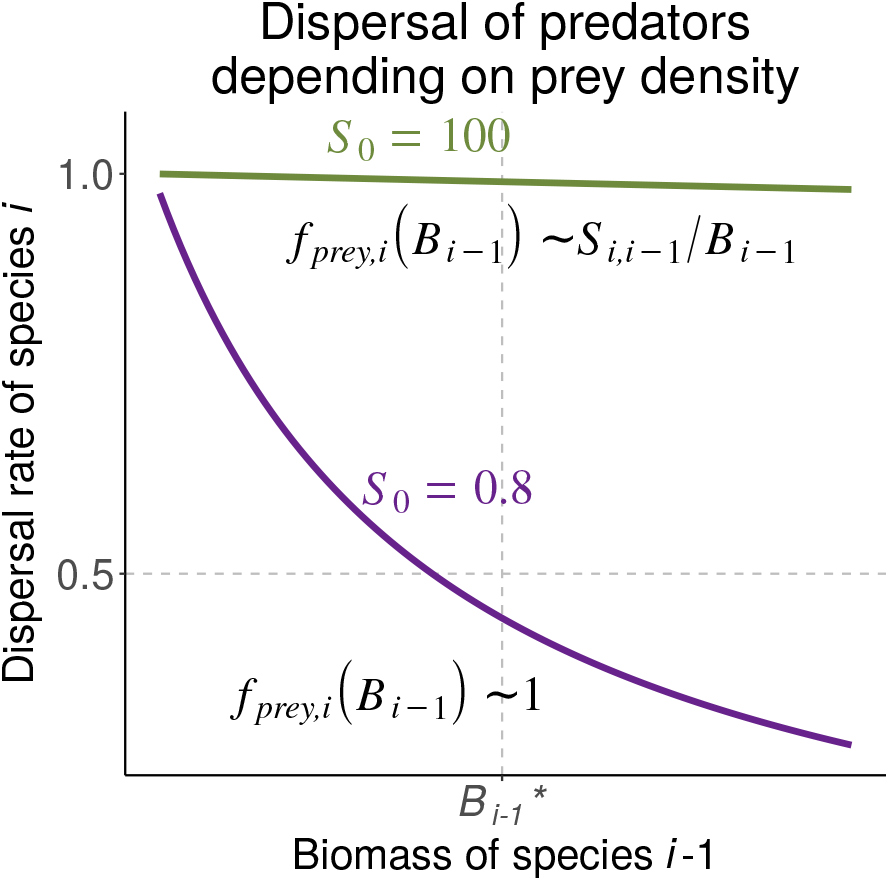
Function describing the dispersal component representing the effect of prey density on predator dispersal: 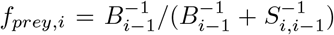 with 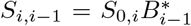. For a sensitivity coefficient *S* = 100 (green), *f*_*prey,i*_ is almost independent of *B*_*i*−1_ in the vicinity of the equilibrium biomass 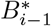 while it is similar to an reverse function of *B*_i−1_ for *S*_0*,i*_ = 100 (purple).

**Figure S2-5:**
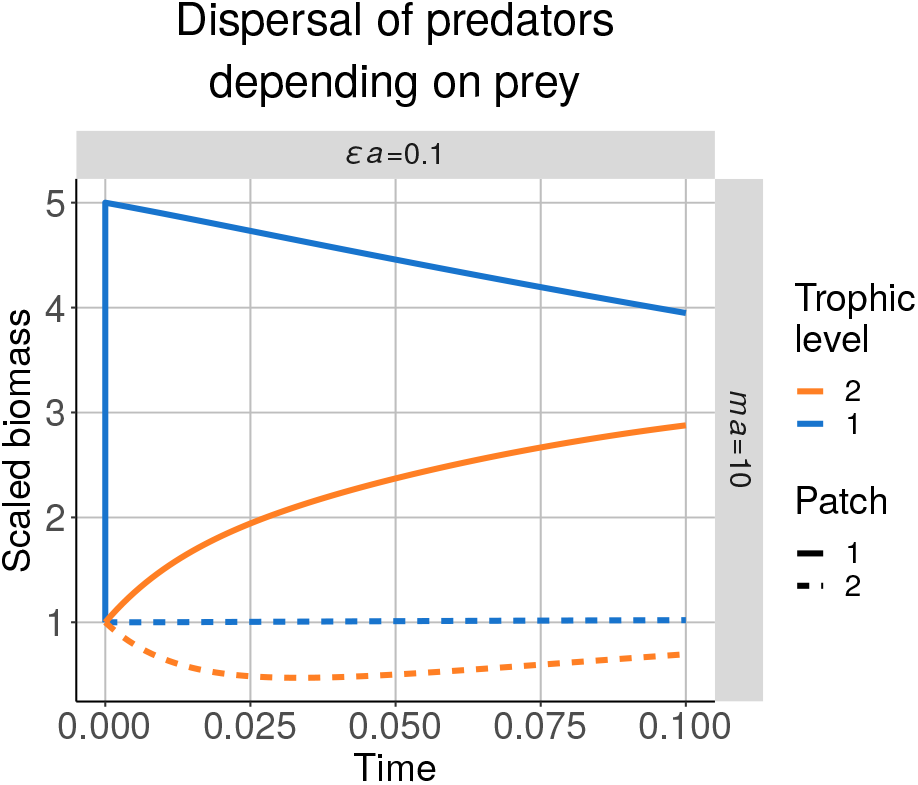
Short-term time series of a metacommunity with two patches and a predator-prey system for a negative effect of predators on prey *ma* = 10 and a positive effect of prey on predators *ϵa* = 0.1. Prey are perturbed (increase of 20% in prey biomass in patch #1 at *t* = 0), and only the predators are able to disperse (*d*_2_ = 1, *S*_0*,i*_ = 10^−3^). Biomasses are rescaled by their value at equilibrium.

**Figure S2-6:**
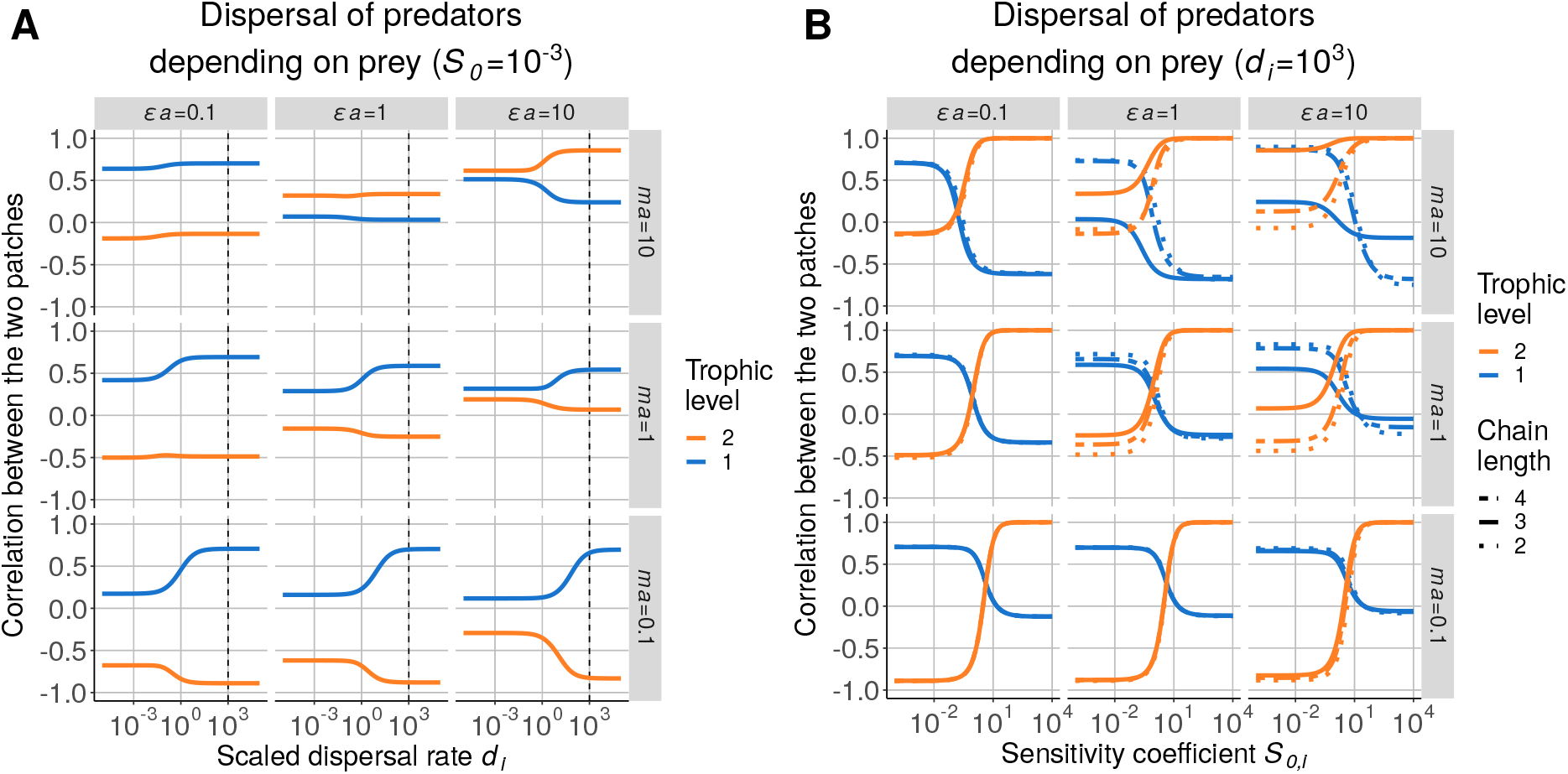
Effect of top-down control and food chain length on the influence of density-dependent dispersal. **A)** Correlation between populations of predators in each patch when dispersal depends on prey biomass for increasing values of scaled dispersal rate *d_i_* (*S*_0*,i*_ = 10^−3^). Prey are perturbed and a top predator (species 3) is present to control the population of species 2. **B)** Correlation between populations in each patch when predator dispersal depends on prey biomass for increasing values of sensitivity coefficient *S*_0*,i*_ (*d_i_* = 10^3^).

### S2-3 Multiple dispersal dependencies

**Figure S2-7:**
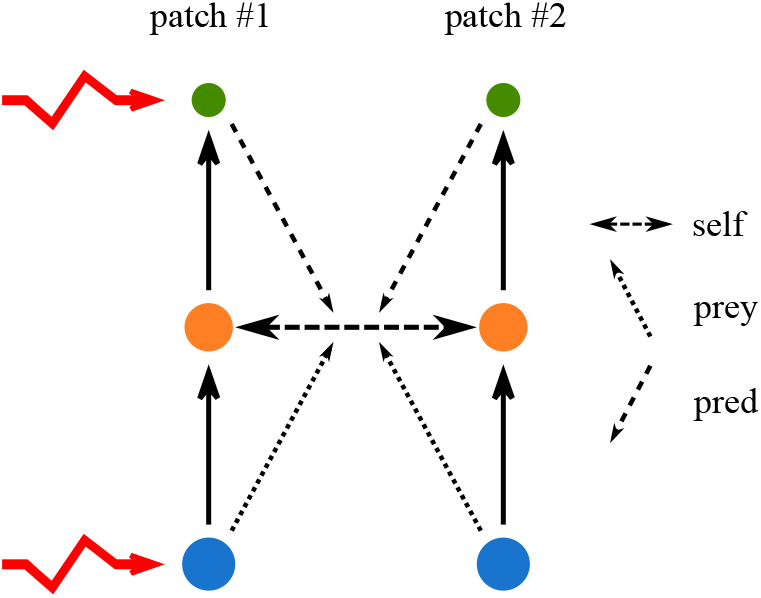
Schema of the metacommunity. Two patches each sustain a food chain with three trophic levels. Primary producers and carnivores in patch #1 experience independent demographic perturbations (*z* = 0.5), and only herbivores are able to disperse. Species dispersal can depend on various combinations of density dependencies: self-dependency, prey dependency and/or predator dependency. Each dependency can be weighted by the effects of self-regulation 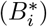, consumption 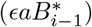 and predation 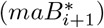 on the growth rate, as detailed in equations (7a-c).

Here, we aim to disentangle the effects of each dependency when several independent perturbations affect the system. Thus, we consider a metacommunity with two patches, each sustaining a food chain with three trophic levels (Fig. S2-7). Only herbivores are able to disperse, and primary producers and carnivores in patch #1 receive independent demographic perturbations (*z* = 0.5, see equation (9) in the main text). Since demographic perturbations maintain a similar ratio of the generated variance to the perturbation variance for all species regardless of their abundance (see Arnoldi et al. (2019)), we can avoid biases while assessing the effects of prey dependency and predator dependency.

Again, the self-dependency of dispersal still leads to the same spatial synchrony as passive dispersal (Fig. S2-8).

Now, we consider two setups: one with self-dependency and prey dependency (Fig. S2-9) and one with self-dependency and predator dependency (Fig. S2-10). These dependencies can also be weighted by local demographic processes (equation (7) and Fig. S2-7).

**Figure S2-8:**
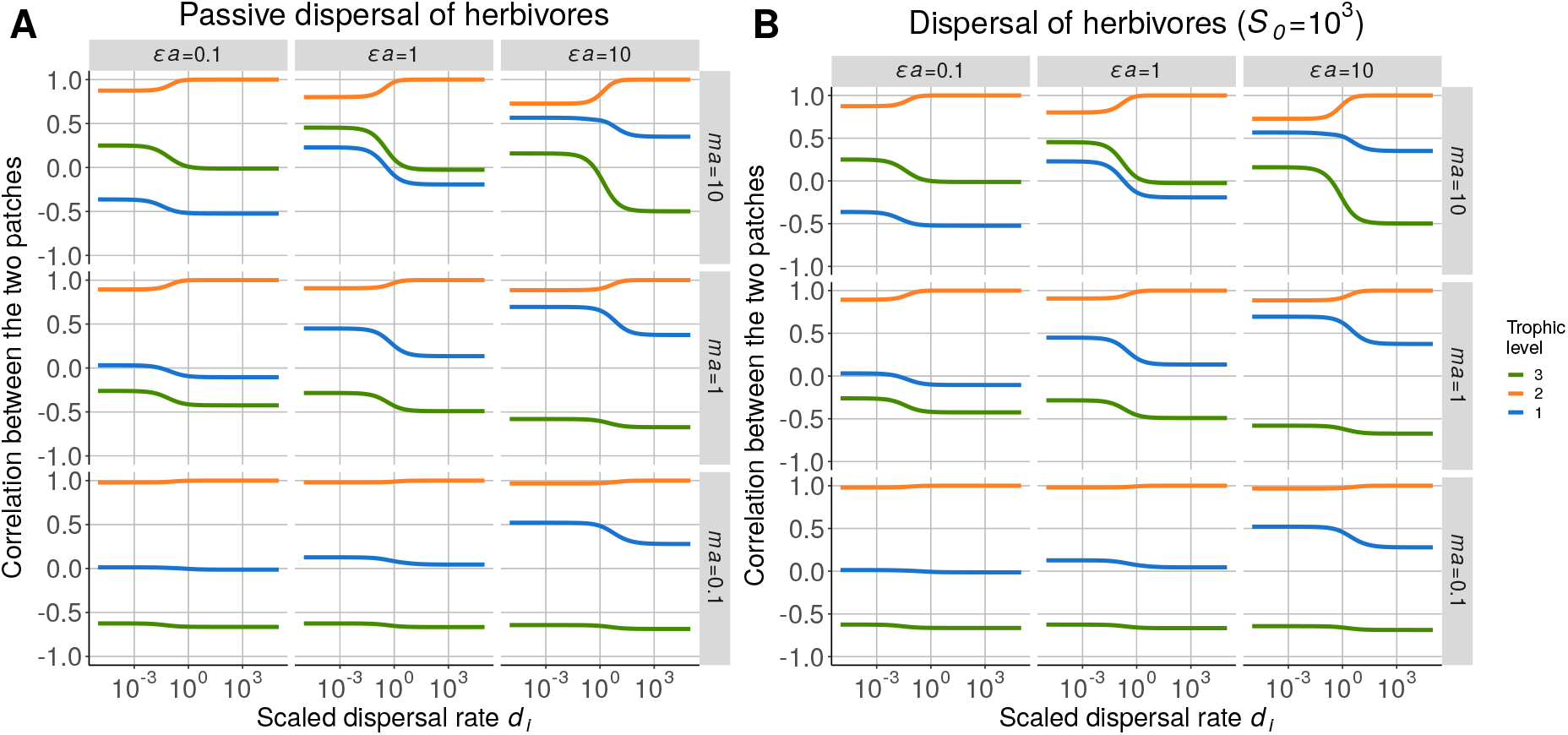
Correlation between populations in each patch in the case of **A)** passive dispersal (classic mass effect) and **B)** dispersal depending on the biomass of the dispersing species (*S_0,i_* = 10^−3^).

**Figure S2-9:**
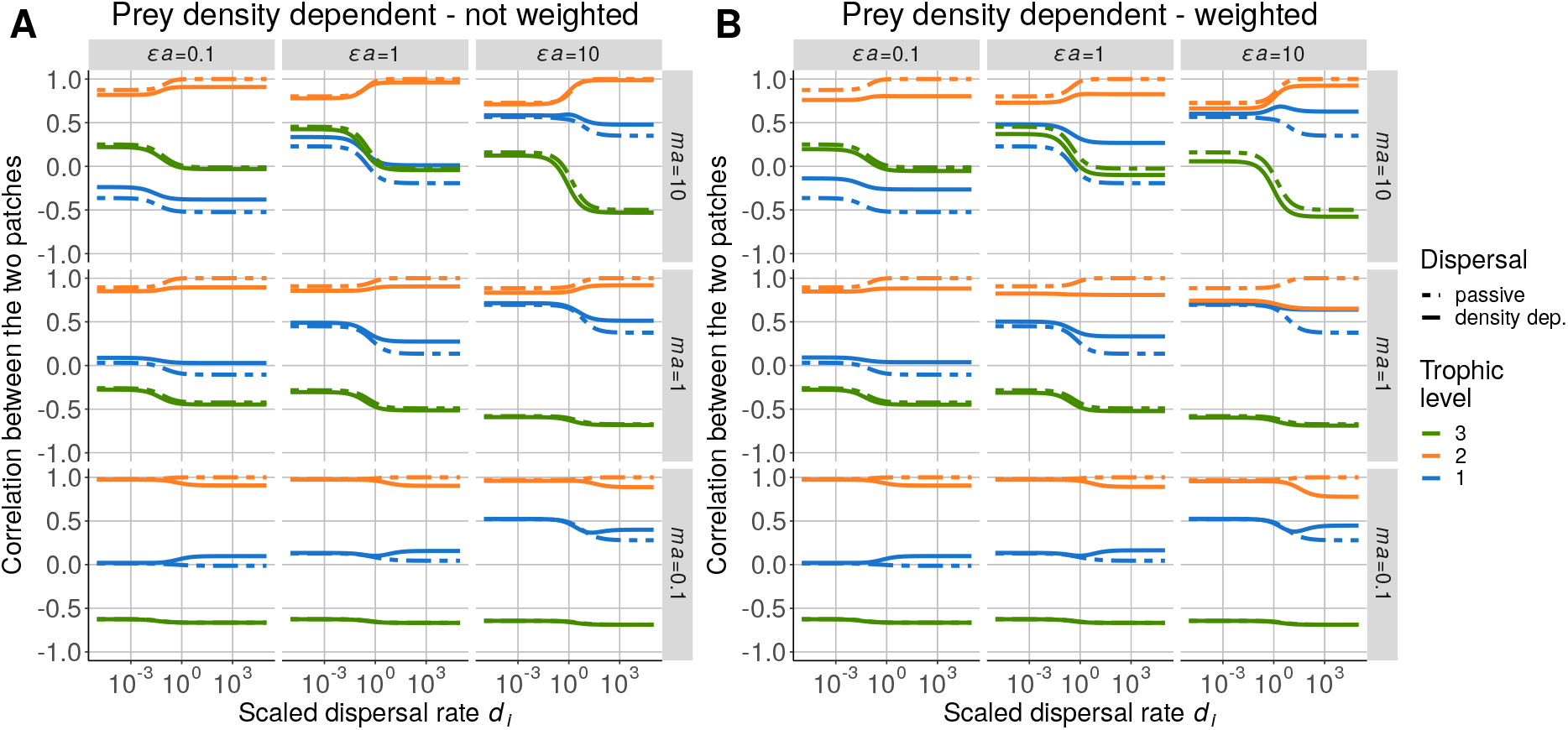
Correlation between populations in each patch when dispersal depends on the biomass of the dispersing species and prey biomass (*S*_0*,self*_ = 10^3^ and *S*_0*,prey*_ = 10^−3^). **A)** Two components of dispersal are not weighted. **B)** Two components of dispersal are weighted.

First, prey dependency leads to a spatial synchrony similar to the case with passive dispersal when dependencies are not weighted (Fig. S2-9A), as herbivores have a higher biomass CV than primary producers (Fig. S2-12C). When dependencies are weighted (Fig. S2-9B), we see small deviations from the model with passive dispersal for high values of positive effect of prey on predators *ϵa* and negative effect of predators on prey *ma*, as prey dependency contributes to more than 50% of the dispersal function of herbivores (species 2) (Fig. S2-11A). These deviations are small because the large contribution of prey dependency is mitigated by the low biomass CV of primary producers compared to herbivores.

Second, predator dependency leads to opposite correlations compared to the model with passive dispersal when dependencies are not weighted (Fig. S2-10A) as carnivores generally have the highest biomass CV (Fig. S2-12C). This does not hold for high values of *ϵa*, as carnivores have a lower (or equivalent) biomass CV than herbivores. When dependencies are weighted, the spatial synchrony strongly changes for low values of *ϵa* and *ma* (Fig. S2-10B) because the low contribution of predator dependency mitigates its strong impact due to the high biomass CV of carnivores. The opposite process acts for high values of *ϵa* and *ma*, while nothing changes for contrasted values of *ϵa* and *ma* (top-left corner and bottom-right corner), as self- and predator-dependencies act in a balanced way.

**Figure S2-10:**
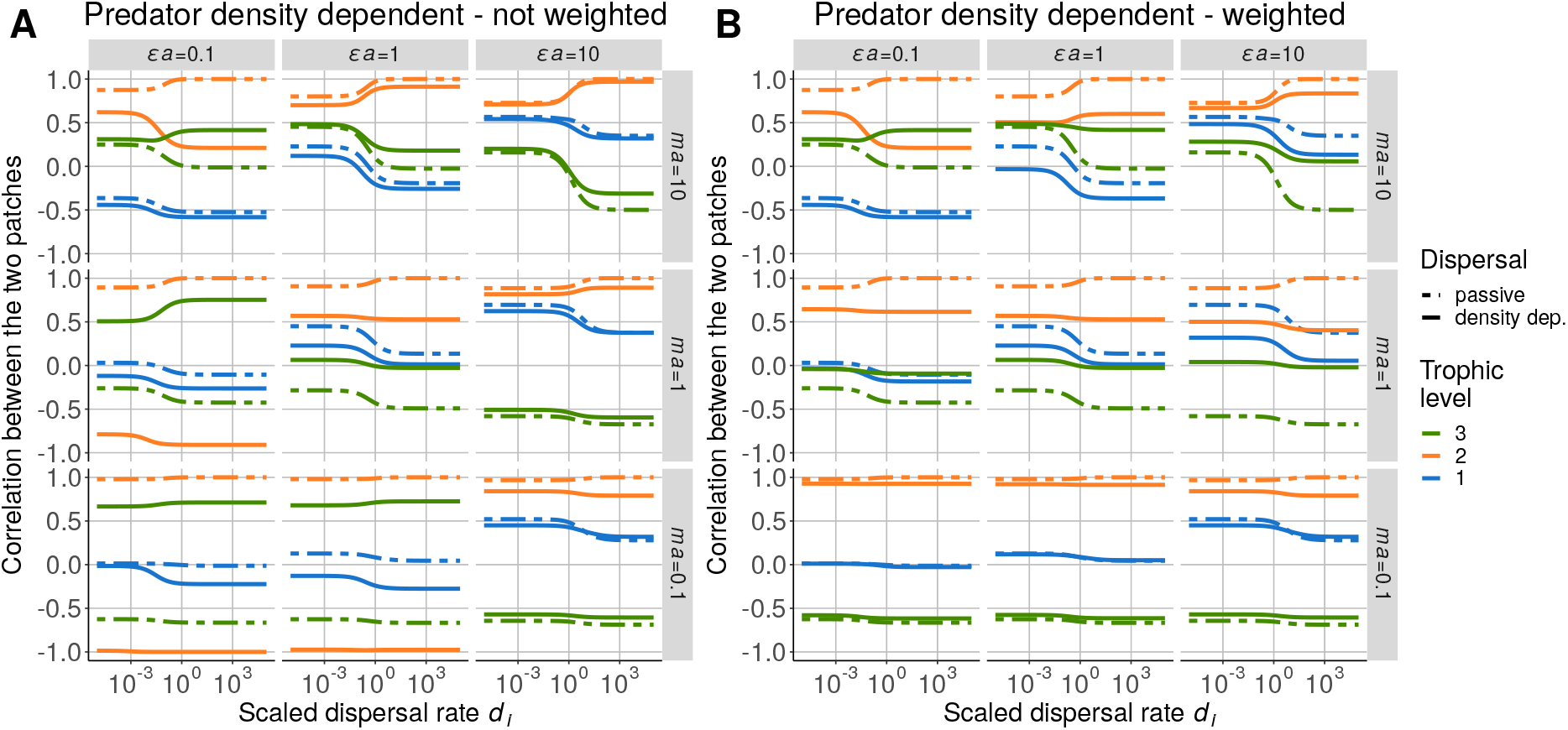
Correlation between populations in each patch when dispersal depends on the focal species and predator abundances (*S*_0*,self*_ = 10^3^ and *S*_0*,pred*_ = 10^3^). Primary producers and carnivores experience independent demographic perturbations, and only herbivores are able to disperse. **A)** Two components of dispersal are not weighted. **B)** Two components of dispersal are weighted by the associated demographic processes.

**Figure S2-11:**
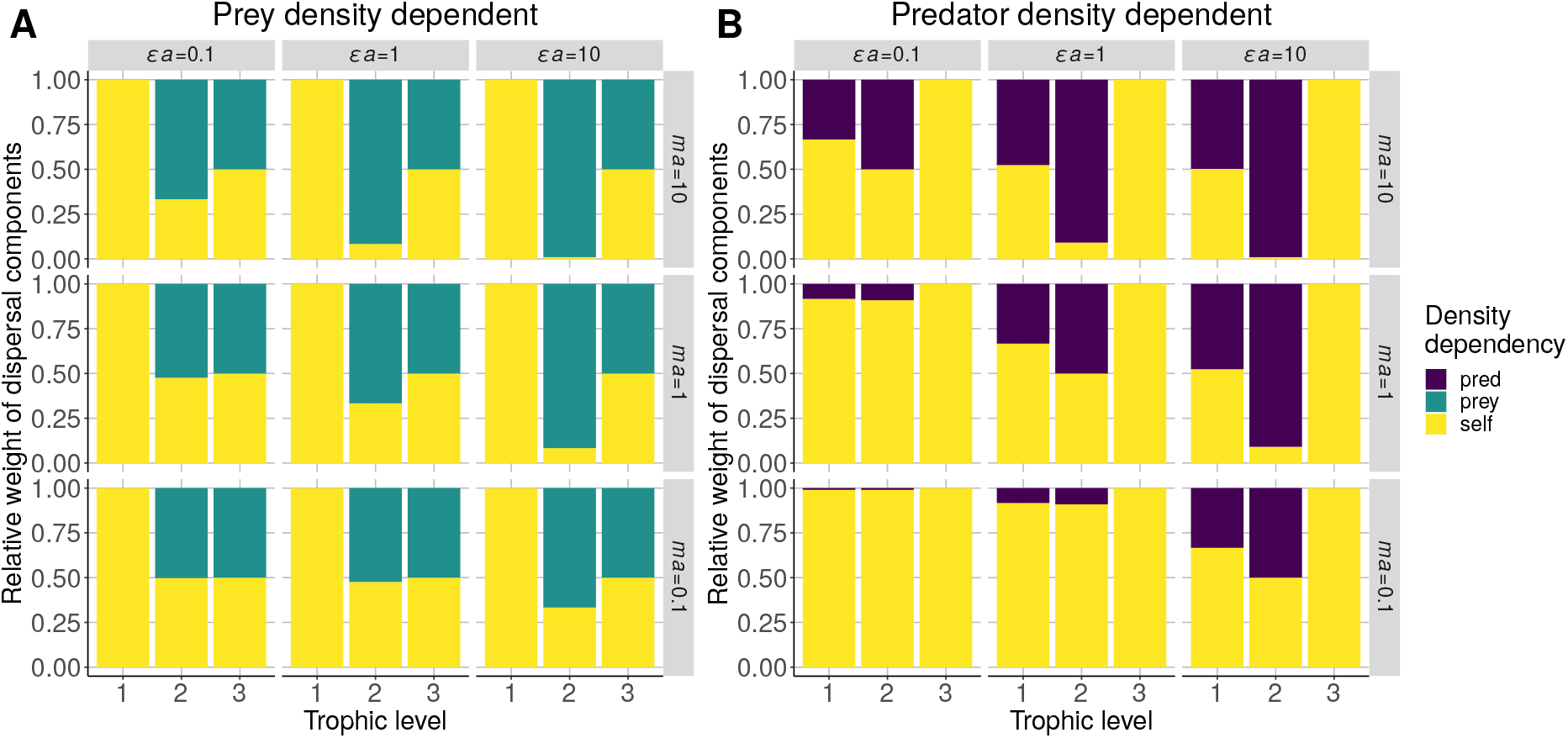
Relative weight of each density dependency in the dispersal function. **A)** Self and prey biomass dependency only. **B)** Self and predator biomass dependency only.

Note that the results in the top-right corner (high values of *ϵa* and *ma*) are subject to changes depending on food chain length.

Now, we understand how prey dependency and predator dependency act when several perturbations affect the system, and we can study the effects of these dependencies together. When dependencies are not weighted (Fig. S2-12A), the spatial synchrony is very similar to the one seen seen in Fig. S2-10A (only self-dependency and predator dependency), as predators have the highest biomass CV in most cases (Fig. S2-12C). When herbivores have the highest biomass CV (top-right corner), correlations are similar to the passive dispersal case (as in Fig. S2-9A).

When dependencies are weighted, the spatial synchrony is similar to the case with passive dispersal for low values of positive effect of prey on predators *ϵa* and negative effect of predators on prey *ma* (Fig. S2-12B) because the low contribution of predator dependency (Fig. S2-12D) mitigates its strong impact due to the high biomass CV of carnivores (Fig. S2-12C). For higher values of *ϵa* and *ma*, we obtain slight changes because prey and predator dependencies still act in a balanced way. Thus, none of them dominates the dispersal function and overrides the effect of biomass CVs.

**Figure S2-12:**
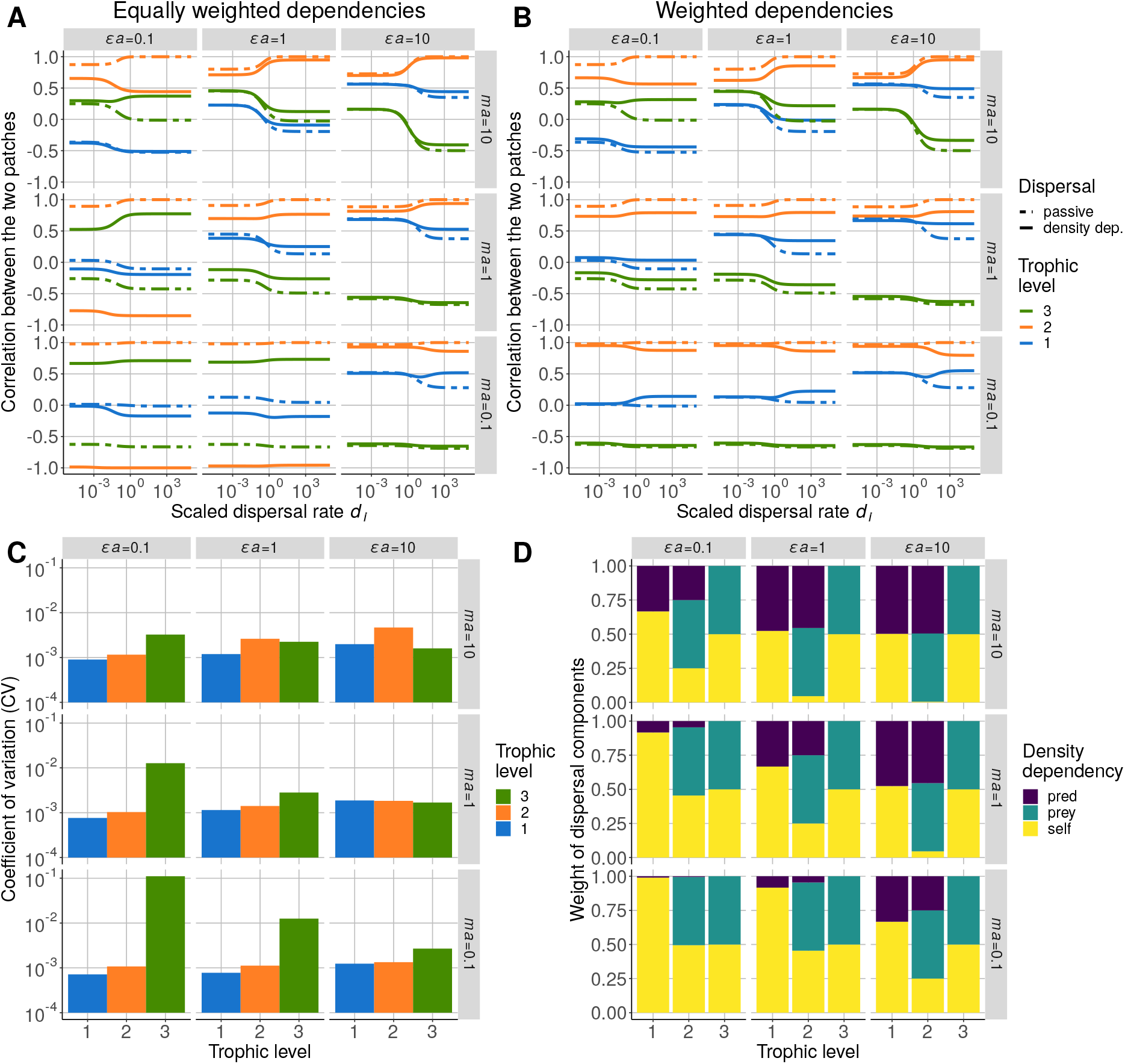
Correlation between populations in each patch when dispersal depends on the focal species, prey and predator abundances (*S*_0*,self*_ = 10^3^, *S*_0*,prey*_ = 10^−3^ and *S*_0*,pred*_ = 10^3^). Primary producers and carnivores experience independent demographic perturbations and only herbivores are able to disperse. **A)** Three components of dispersal are not weighted. **B)** Three components of dispersal are weighted. **C)** Biomass CV of each species. **D)**Weight of each density dependency in the dispersal function.

### S2-4 Environmental perturbation and multiple dispersal dependencies

**Figure S2-13:**
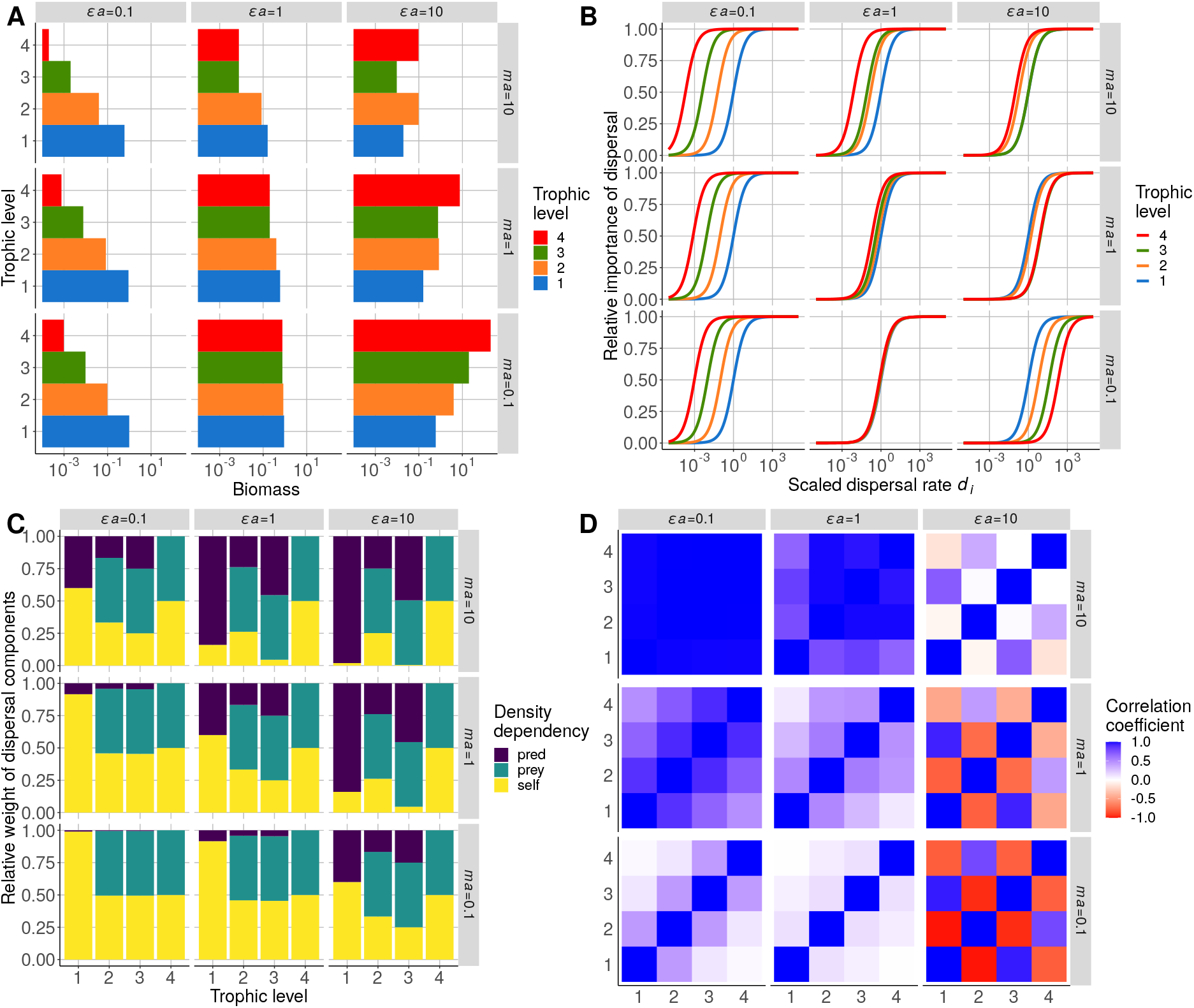
Response of a four-species food chain to correlated environmental perturbations affecting all species. **A**) Biomass distribution. **B**) Relative importance of dispersal processes compared to local demographic processes (see Box 3 in the main text). **C**) Relative weight of each density dependency in the dispersal function *F_i_* (*B_i-1_, B_i_, B_i+1_*). **D)** Correlation between populations in each patch in the case of passive dispersal. All species disperse at the same scaled rate *di* and all species in patch #1 experience correlated environmental perturbations.

The spatial synchrony in Fig. 6A in the main text is due to environmental perturbations that affect more abundant species (*i.e.*, the ratio of the generated variance to the variance of perturbations is higher for abundant species (see Arnoldi et al., 2019 for more details)), which are primary producers (Fig. S2-13A) and lead to a bottom-up propagation of perturbations in patch #1 correlating all its species (Fig. S2-13D). Because dispersal affects the less abundant species the most (Fig. 7A), perturbations are mainly transmitted by higher trophic levels for a positive effect of prey on predators *ϵa* = 0.1 and a negative effect of predators on prey *ma* = 10 (Fig. 6 in the main text), thus leading to a top-down transmission of perturbations in patch #2, which anticorrelates adjacent trophic levels. These two different correlation patterns in patches #1 and #2 lead to the correlation or anticorrelation of the two populations of each species as described in Box 1 and detailed in Quévreux et al. (2021).

Even if dispersal has a low relative importance in the overall dynamics of carnivores (Fig. S2-13B) for *ϵa* = 0.1 and *ma* = 10, it strongly affects the spatial synchrony. In fact, at low scald dispersal rate *d_i_*, the spatial synchrony obtained in Fig. S2-18A (only top predators are able to disperse) does not correspond to Fig. 6B, while Fig. S2-18B (top predators and carnivores are able to disperse) successfully reproduces the spatial synchrony.

**Figure S2-14:**
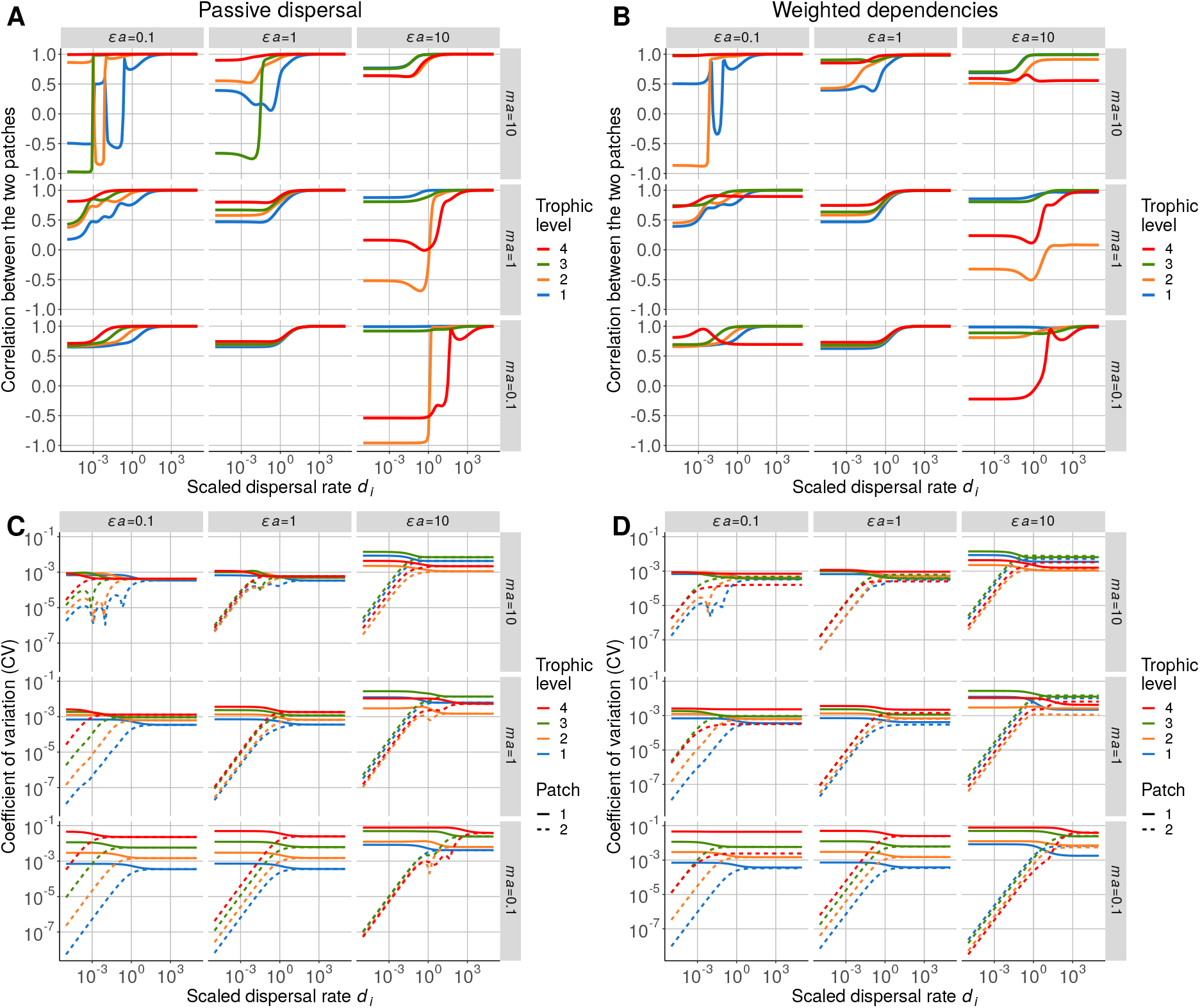
Response of a metacommunity where all species in patch #1 experience correlated environmental perturbations and are all able to disperse. Correlation between populations in each patch when dispersal is **A)** passive or **B)** depends on self, prey and predator biomass (*S*_0*,self*_ = 10^3^, *S*_0*,prey*_ = 10^−3^ and *S*_0*,pred*_ = 10^3^) weighted by their contribution to the growth rate. Biomass CV of each species in each patch when dispersal is **C)** passive or **C)** density-dependent with weighted contributions.

In Fig. S2-15, we detail the dynamics of the metacommunity after a pulse perturbation to explain the spatial synchrony observed in Fig. 6 in the main text. Dispersal is low (*d_i_* = 10^−4^) and mainly matters for top predators (Fig. S2-13B), who transmit perturbations from patch #1 to patch #2. In the case of passive dispersal (Fig. S2-15A), dispersal balances the biomass of top predators between the two patches, thus leading to a decrease in patch #1 right after the perturbation and an increase in patch #2. This leads to an increase in the biomass of carnivores in patch #1 due to the abundance of prey and the rapid decrease in predator while it leads to a decrease in patch #2 due to the increase in top predator biomass. Therefore, the two populations of carnivores are anticorrelated.

In the case of density-dependant dispersal (Fig. S2-15B), prey density-dependent dispersal first amplifies the effect of the pulse perturbation because top predators migrate more to patch #1 because of prey abundance. This leads to an increase in the biomass of carnivores in patch #2 because predator density-dependent dispersal that makes carnivores to migrate to patch #2 where top-predator biomass is decreasing. Then, the biomass of top predators decreases in patch #1 to converge to its equilibrium value and it also decreases in patch #2 because of prey density-dependent dispersal stall has a strong effect because of the persisting unbalance in prey biomass, thus correlation the populations of top predators. Finally, the biomass of carnivores in both patches decreases to converge to the equilibrium, thus correlating the two populations of carnivores.

**Figure S2-15:**
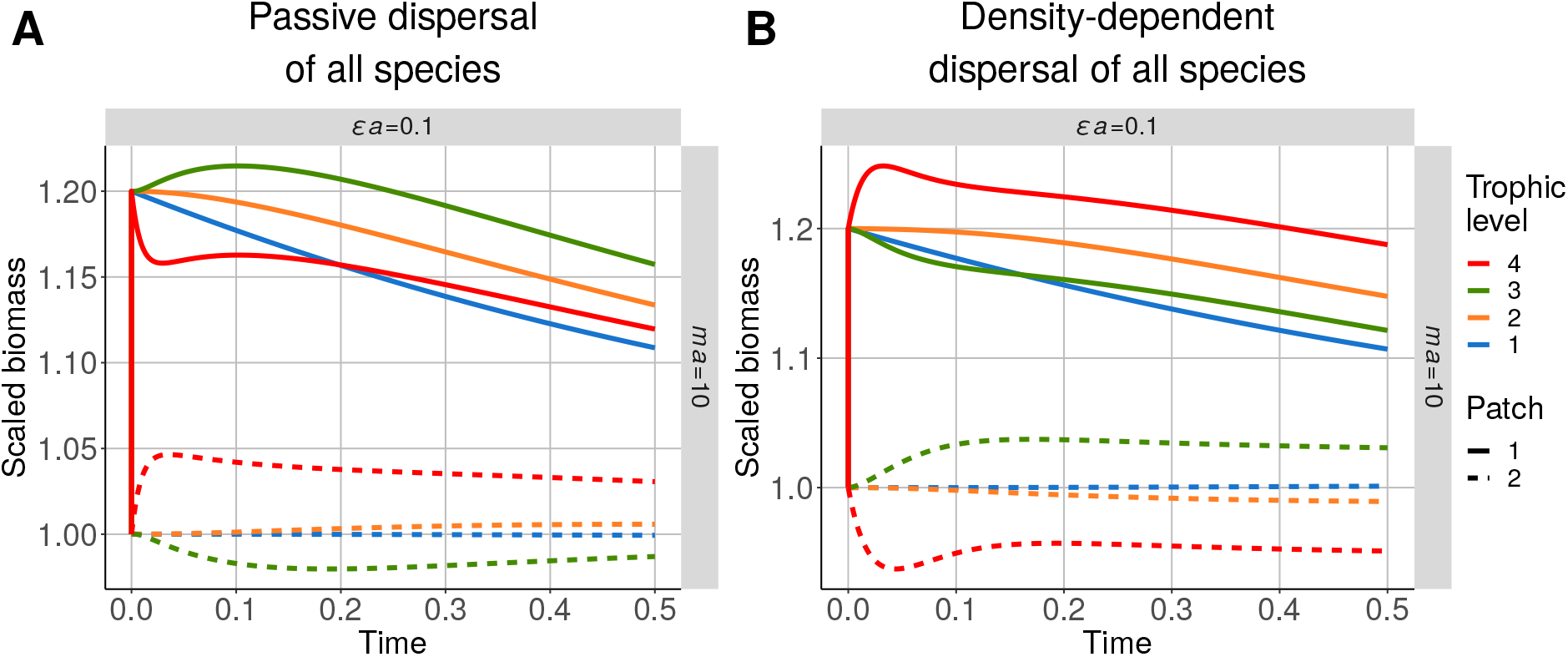
Short-term time series of a metacommunity with two patches and a predator-prey system for *ma* = 10 and *ϵa* = 0.1. All species are perturbed (increase of 20% in prey biomass in patch #1 at *t* = 0), and all species are able to disperse (*d_i_* = 10^−4^). **A)** Passive dispersal. **B)** Dispersal depends on self, prey and predator biomass (*S*_0*,self*_ = 10^3^, *S*_0*,prey*_ = 10^−3^ and *S*_0*,pred*_ = 10^3^) weighted by their contribution to the growth rate. Biomasses are rescaled by their value at equilibrium.

**Figure S2-16:**
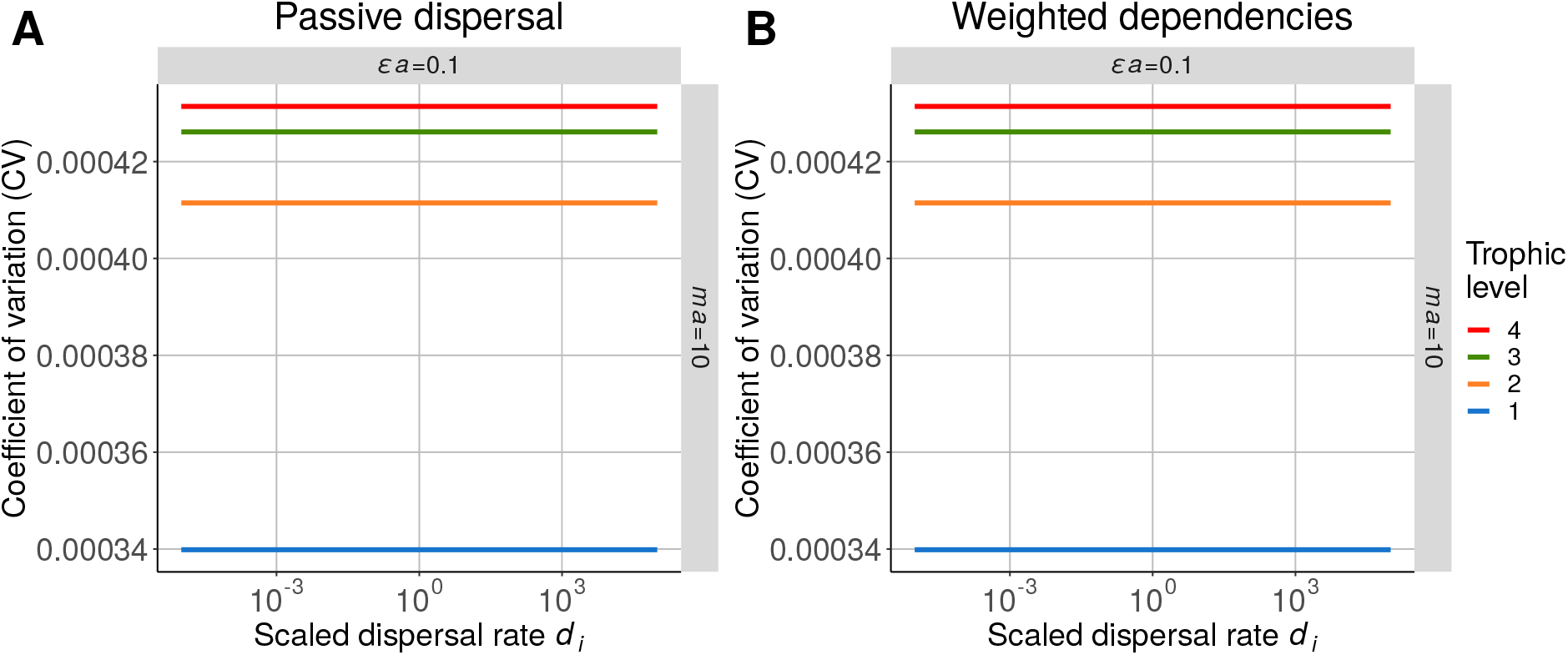
Coefficient of variation (CV) of the entire population of each species at the metacommunity scale. All species in patch #1 experience correlated environmental perturbations (positive effect of prey on predators *ϵa* = 0.1 and negative effect of predators on prey *ma* = 10). Dispersal can be **A)** passive or B) dependent on the self, prey and predator biomass (*S*_0*,self*_ = 10^3^, *S*_0*,prey*_ = 10^−3^ and *S*_0*,pred*_ = 10^3^) weighted by their contribution to the growth rate.

**Figure S2-17:**
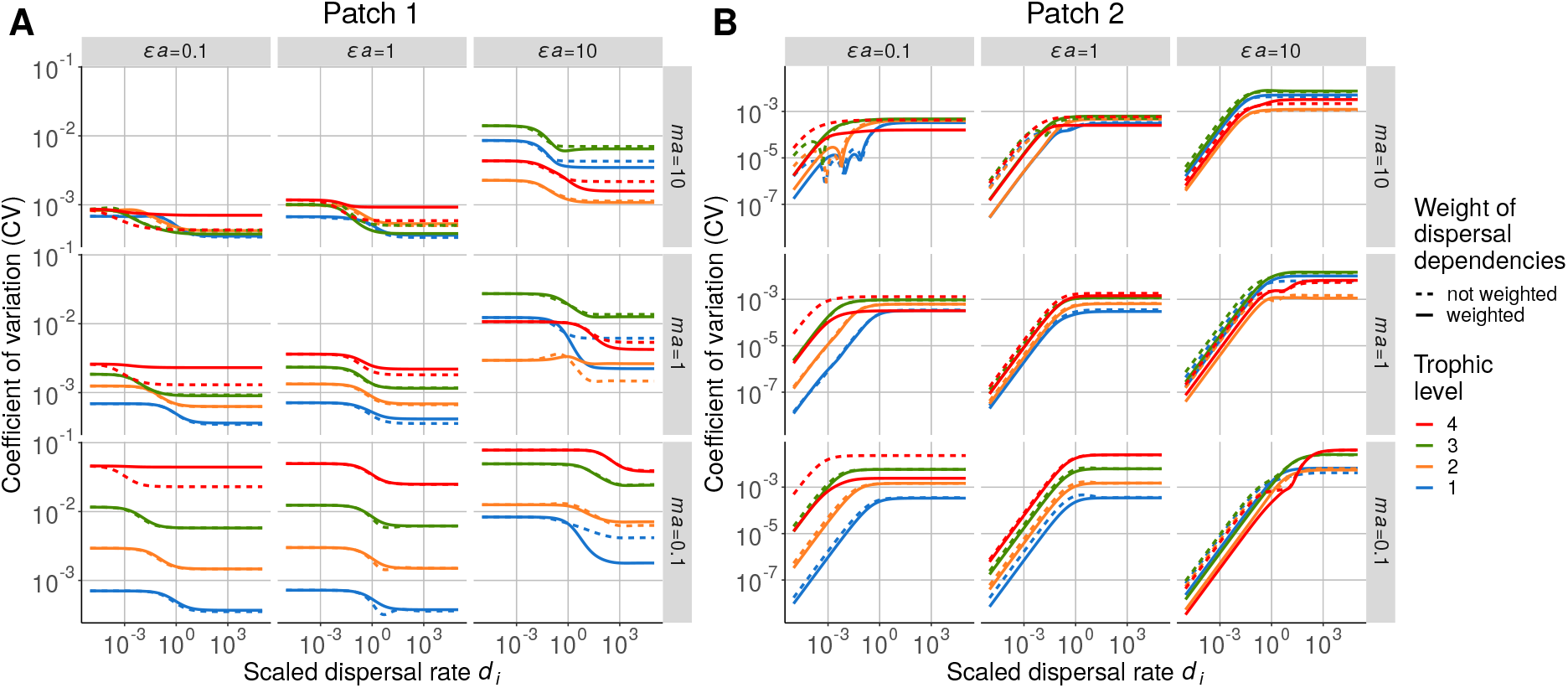
Coefficient of variation (CV) of each species in **A)** patch #1 and **B)** patch #2. All species in patch #1 experience correlated environmental perturbations (positive effect of prey on predators *ϵa* = 0.1 and negative effect of predators on prey *ma* = 10). Dispersal depends on self, prey and predator biomass (*S*_0*,self*_ = 10^3^, *S*_0*,prey*_ = 10^−3^ and *S*_0*,pred*_ = 10^3^) weighted (solid line) or not (dashed line) by their contribution to the growth rate.

**Figure S2-18:**
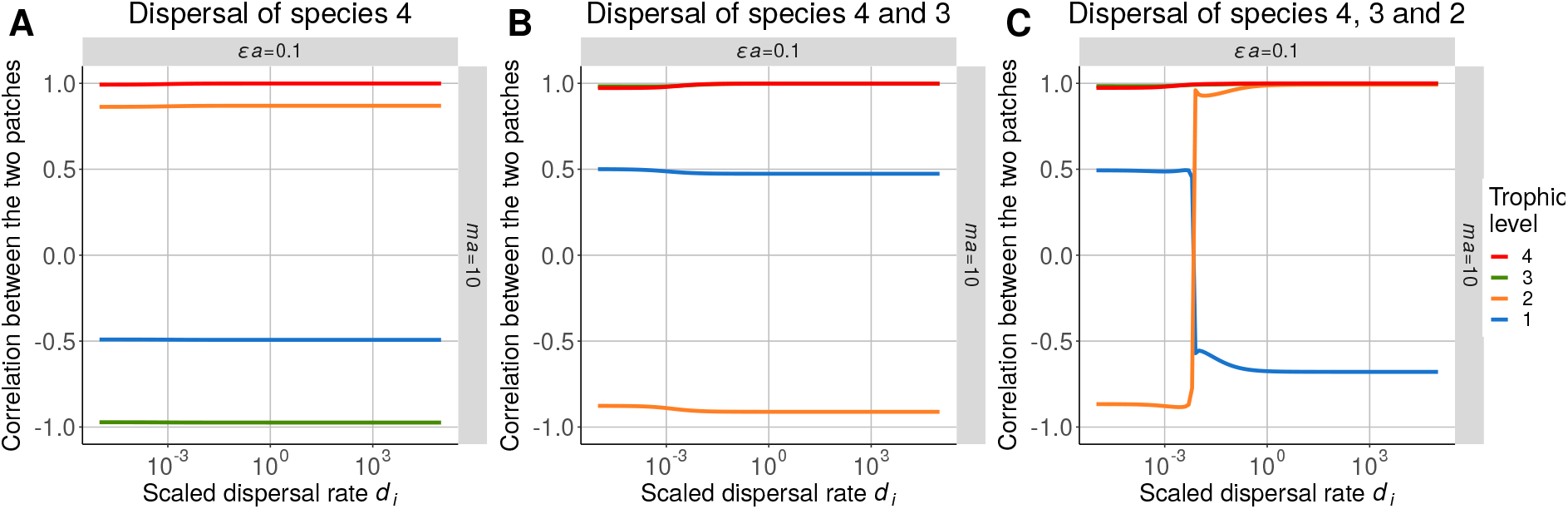
Correlation between populations in each patch when dispersal depends on self, prey and predator biomass (*S*_0*,self*_ = 10^3^, *S*_0*,prey*_ = 10^−3^ and *S*_0*,pred*_ = 10^3^) weighted by their contribution to the growth rate. All species in patch #1 experience correlated environmental perturbations, and **A)** only top predators, **B)** top predators and carnivores or **C)** all species except primary producers are able to disperse (positive effect of prey on predators *ϵa* = 0.1 and negative effect of predators on prey *ma* = 10).

### S2-5 Linear density-dependent functions

The density-dependent function *F_i_*(*B_i−1_, B_i_*, *B*_*i*+1_) can also be defined by linear components:

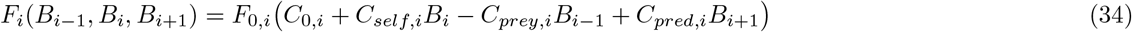

The weights of each component are now defined as follows:

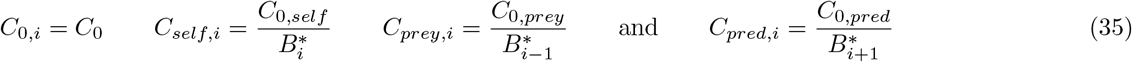

The spatial synchrony obtained in Fig. S2-19A, S2-19B and S2-19C are extremely similar to those in Fig. S2-2A, 3B and 3A, respectively. However, the prey-dependent component is a negative function. Because *F_i_*(*B_i-1_, B_i_, B_i+1_*) must be positive to ensure consistent migration flows, the term *C*_0*,i*_ - *C_prey,i_B_i-1_* must be positive. If *C*_0*,i*_ is small, then the term is negative and the model is wrong. If *C*_0*,i*_ is large, then the prey dependency becomes negligible (Fig. S2-19D). This finding is consistent with the results from Fig. 3C. Therefore, the functions defined by Hauzy et al. (2010) and used in the main text are better because they are positive and symmetric.

**Figure S2-19:**
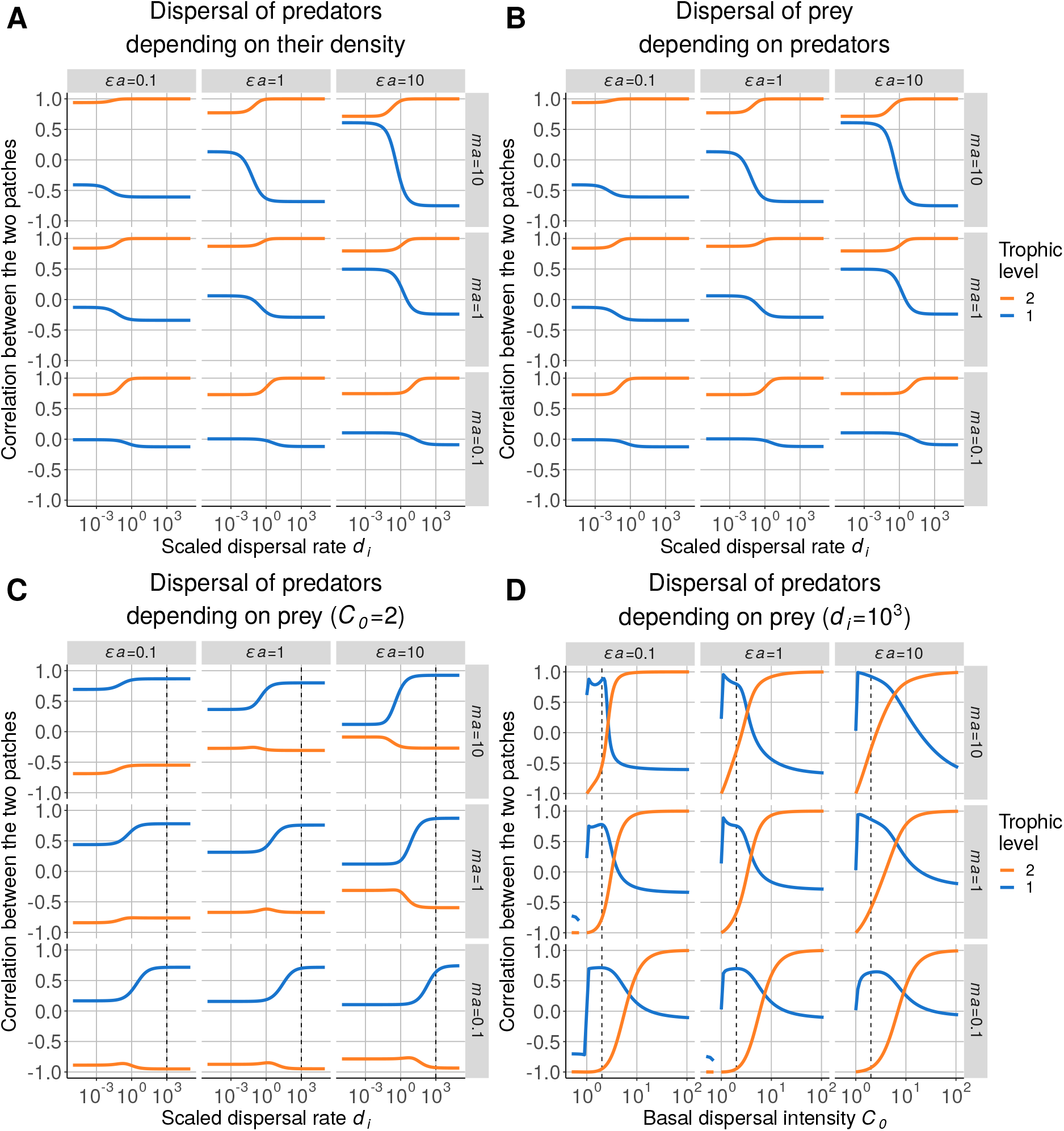
Correlation between the populations in each patch when the dispersal components are linear functions of species biomasses. **A)** Dispersal of predators depending on their own density (*C*_0*,self*_ = 1), with prey experiencing demographic perturbations in patch #1. **B)** Dispersal of prey depending on predator density (*C*_0*,pred*_ = 1), with predators experiencing demographic perturbations in patch #1. **C)** Dispersal of predators depending on their own density (*C*_0_ = 2 and *C*_0*,prey*_ = 1), with prey experiencing demographic perturbations in patch #1. **D)** Effect of passive dispersal intensity *C*_0_ on prey-dependent dispersal of predators (*C*_0*,prey*_ = 1 and *d_i_* = 10^3^). The dashed lines represent the values of *d_i_* and *C*_0_ between graphs **C** and **D**.

